# Pairs of amino acids at the P- and A-sites of the ribosome predictably and causally modulate translation-elongation rates

**DOI:** 10.1101/2020.04.20.051839

**Authors:** Nabeel Ahmed, Ulrike A. Friedrich, Pietro Sormanni, Prajwal Ciryam, Naomi S. Altman, Bernd Bukau, Günter Kramer, Edward P. O’Brien

## Abstract

Variation in translation-elongation kinetics along a transcript’s coding sequence plays an important role in the maintenance of cellular protein homeostasis by regulating co-translational protein folding, localization, and maturation. Translation-elongation speed is influenced by molecular factors within mRNA and protein sequences. For example, the presence of proline in the ribosome’s P- or A-site slows down translation, but the effect of other pairs of amino acids, in the context of all 400 possible pairs, has not been characterized. Here, we study *Saccharomyces cerevisiae* using a combination of bioinformatics, mutational experiments, and evolutionary analyses, and show that many different pairs of amino acids and their associated tRNA molecules predictably and causally encode translation rate information when these pairs are present in the A- and P-sites of the ribosome independent of other factors known to influence translation speed including mRNA structure, wobble base pairing, tripeptide motifs, positively charged upstream nascent chain residues, and cognate tRNA concentration. The fast-translating pairs of amino acids that we identify are enriched four-fold relative to the slow-translating pairs across *Saccharomyces cerevisiae*’s proteome, while the slow-translating pairs are enriched downstream of domain boundaries. Thus, the chemical identity of amino acid pairs contributes to variability in translation rates, elongation kinetics are causally encoded in the primary structure of proteins, and signatures of evolutionary selection indicate their potential role in co-translational processes.

## INTRODUCTION

The rates associated with translation determine the time scales of protein synthesis [1,2], influence protein expression levels [3], and can affect the structure and function of the protein produced [4–6]. While the rate of translation initiation is a key kinetic parameter influencing protein expression levels, the non-uniform rate of translation elongation across coding sequences can influence the fate of nascent proteins and the downstream cellular processes they take part in. Variation in translation-elongation kinetics influences protein homeostasis by modulating co-translational protein folding, localization, and maturation [6–8]. Hence, characterizing the molecular factors that determine the rate at which individual codon positions along a transcript are translated aids our understanding of how a functional proteome is regulated and produced *in vivo*.

The rate of translation elongation was originally thought to be determined only by the A-site codon’s cognate tRNA concentration as it influences the rate of tRNA accommodation into the A-site [9,10]. More frequently used codons across a transcriptome were presumed to be translated at faster rates as they typically have a higher abundance of cognate tRNAs. Over the past decade some ten other molecular factors have been identified that can influence the rate of translation elongation [7,11] including features inherent to the mRNA sequence, such as mRNA structure [12–14], and features inherent to the protein sequence, such as the presence of particular tripeptide sequence motifs composed of one or more prolines [15–19] and positively charged nascent-chain residues within the negatively charged ribosome exit tunnel [20–22]. There are still unexplored molecular factors that have the potential to influence translation elongation rates. Furthermore, the aforementioned protein-based factors suggest the intriguing possibility that the primary structures of proteins, beyond proline containing motifs [15], have the potential to causally encode translation-elongation rate information.

Since the ribosome catalyzes peptide bond formation between 400 unique amino acid pairs when they reside in the P- and A-sites of the ribosome, we hypothesized that the chemical identity of the P-site amino-acid, in the context of these pairs, could influence translation speed at the A-site in a predictable and causal way, beyond known effects arising from proline-containing pairs and cognate tRNA concentration. In this study, we utilize ribosome profiling data generated from *Saccharomyces cerevisiae* to test this hypothesis. We bioinformatically isolated the effect of the P-site amino acid and tRNA identities on translation at the A-site by keeping the A-site amino acid fixed, as this controls for cognate tRNA concentration and accommodation kinetics. We identified certain amino acids, that when present in the P-site, appear to either speed up or slow down the rate of translation when a particular amino acid is present in the A-site. We experimentally tested these predictions by mutating the P-site amino acid and detected the change in ribosome density, which is a function of translation speed, via ribosome profiling. While an amino acid effect has been well established for proline, this is the first study to identify a large number of amino acid pairs for which the chemical identity of the P-site amino acid and tRNA systematically influences the translation elongation rate at the A-site. Finally, we demonstrate that across the *Saccharomyces cerevisiae* proteome there are signatures of evolutionary selection pressure on the fast- and slow-translating amino acid pairs we have identified that suggests these pairs might play a role in regulating the co-translational maturation of proteins.

## RESULTS

### Beyond proline, the identity of the P-site amino acid can influence the translation rate at the A-site

Ribosome profiling is a high-throughput technique that measures ribosome densities that are a function of the location and number of ribosomes translating different codon positions across a transcriptome [23]. The measured normalized ribosome density *ρ* at a codon position is equal to the number of reads mapped to that position divided by the average number of reads per codon in the coding sequence in which that codon resides. *ρ* at a codon is inversely related to the speed at which ribosomes translate that codon [24]. A *ρ* greater than 1.0 indicates that there is slower than average translation elongation rate while a *ρ* less than 1.0 reflects faster than average translation. Thus, from one codon position to the next along a transcript, a ribosome can be said to speed-up or slow-down its elongation speed reflected in the variation of *ρ*.

We analyzed the translational profiles of 364 high-coverage transcripts (Data S1) measured in six independent, published data sets from four different labs [25–29] (Table S1). These datasets where chosen because they have high read coverage and do not use cycloheximide (CHX) pre-treatment, as CHX has been shown to artificially distort ribosome profiles [30]. The 364 transcripts were chosen after applying a filter of at least 3 reads per codon in the dataset with the highest coverage [26]. This high coverage filter allows us to minimize any sequencing noise and provides a more precise reporter of translation speed. This subset of transcripts is representative of the entire transcriptome as the sequence properties of these transcripts are similarly distributed (see Fig. S8 of ref. 14).

To isolate the effect of the P-site on translation at the A-site we compare the ribosome density when a particular amino acid is present in the P-site versus when it is not. Specifically, for each of the 400 unique pairs of amino acids that can reside in the P- and A-sites — which for a given pair we denote as (*X, Z*), where *X* is the amino acid in the P-site and *Z* is the amino acid in the A-site — we first determined the normalized ribosome density distribution, [*ρ*(*X, Z*)], arising from all instances of the pair (*X, Z*) in the data set as well as the distribution [*ρ*(∼*X, Z*)] arising from all instances of *Z* being in the A-site but *X* not being present in the P-site. For example, for the pair denoted (N, R), N is in the P-site and R is in the A-site, while (∼N, R) corresponds to the 19 other naturally occurring amino acids that can be in the P-site when R is in the A-site (Fig. 1a). We then calculated the percent change (Eq. 1) in the median of [*ρ*(*X, Z*)] relative to the median of [*ρ*(∼*X, Z*)], as this measures whether the identity of the P-site amino acid tends to lead to faster or slower translation relative to when any other amino acid is present in the P-site.

**Figure 1.**
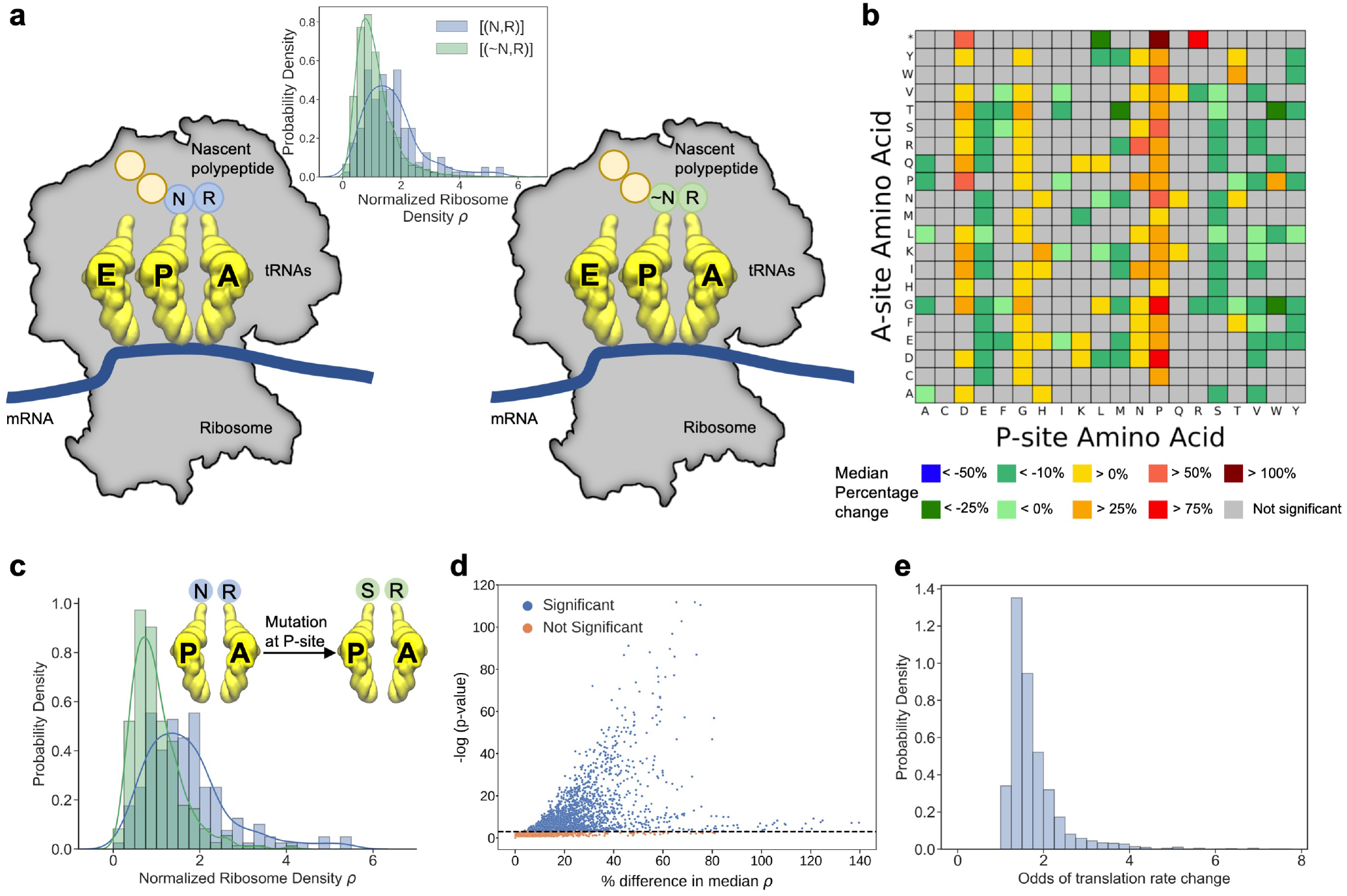
Bioinformatic analyses of ribosome profiling data indicate that the identity of amino acids in the P- and A-sites can predictably alter the translation speed of the A-site codon. **(a)** A ribosome with the amino acids N and R in the P- and A-sites, respectively. From ribosome profiling data, we calculated the distribution of ribosome densities in the A-site from all instances of (N, R) in our dataset and compared the result to the distributions of all other instances of R in the A-site when N is not present in the P-site, denoted [(∼N, R)] (top panel). **(b)** Each box in the matrix indicates, for a pair of amino acids in the P- and A-sites, the percent change in median normalized ribosome density ρ when that particular amino acid is in P-site compared to any other amino acid in the P-site, keeping the A-site amino acid unchanged (Eq. 1). The sign of the percent change must be consistent in all 6 analyzed ribosome profiling datasets and statistically significant in at least 4 out of the 6 datasets, otherwise the box is colored gray. * corresponds to any of the stop codons being present in the A-site. **(c)** Comparison of distributions of amino acid pairs where R is kept constant at the A-site while the P-site is mutated from N to S. The distributions of normalized ribosome densities for P- and A-site pairs (N, R) and (S, R), which differ significantly from each other, are shown (Mann-Whitney U test, *p* = 4.45 × 10^−17^). The median normalized ribosome densities of the two distributions differ by 53.4%, and the odds of a change in translation speed when (N, R) is mutated to (S, R) or vice versa is 2.98 (Eq. S3). **(d)** The estimated percent difference values for all 7,980 mutations of amino acid pairs with a constant A-site are plotted with respect to the statistical significance of the difference between the distributions (see Methods). We estimate that mutating the P-site will lead to significant changes in translation speed in 4,254 (53%) of these mutations. **(e)** For the significant combinations of amino acids pairs, the distribution of the odds of mutating any instance of the pair resulting in a change in speed is plotted.

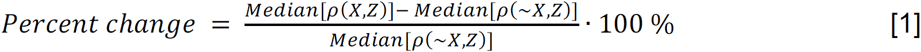

We use the median of these [*ρ*(*X, Z*)] distributions, as opposed to the average, because the median is less sensitive to outlier, long-tail effects in right-skewed distributions typically found in ribosome profiles (Fig. 1a (inset) and 1c). Critically, this analysis approach controls for confounding cognate tRNA concentration effects and accommodation kinetics at the A-site because the A-site amino acid is held fixed. We do this for all possible 20 amino acids in the P-site for a fixed amino acid in the A-site (Fig. S1a). For an amino acid in the P-site, there are 21 possible pairs when we include the presence of stop codons in the A-site (Fig. S1b). Applying this analysis to each of the six published datasets we obtained six, 21-by-20 matrices reporting the percent change (Eq. 1) in ribosome density when a particular amino acid pair is present in the P-and A-sites (Fig. S2).

Given the well-known variability in ribosome profiling results between different experiments [31] we focus on highly robust, reproducible results by (*i*) only drawing conclusions from the sign of the percent change, which indicates either a speed up or slow down between different pairs of amino acids, and by (*ii*) taking the intersection of those pairs that exhibit a consistent sign change and have a percent change that is statistically different than zero in at least four of the datasets. We find 86 pairs in which the presence of a particular P-site amino acid is associated with faster translation (green-shifted colors in Fig. 1b) and 81 pairs in which the identity of the P-site amino acid is associated with slower translation (red-shifted colors in Fig. 1b). The results for the remaining pairs are not significant or are not consistent across the datasets (gray boxes in Fig. 1b). An important, naturally occurring internal control in these data is proline, which is known from biochemical studies [16,32] to tend to slow down translation when present in the P-site. Consistent with those findings, we observe a vertical strip of warm colors in Fig. 1b when Pro is in the P-site, indicating that when paired with almost any other amino acid in the A-site, proline tends to increase ribosome density, *i*.*e*., slow translation down relative to any other amino acid being in the P-site. These results suggest that the identity of the P-site amino acid in 167 pairs of amino acids can, on average, predictably speed up or slow down translation relative to the median speed when any other amino acid is in the P-site.

### Factors known to modulate translation speed do not explain these results

To test whether the potentially confounding factors of tripeptide motifs [15], positively charged upstream residues [20,21], downstream mRNA structure [13,33], cognate tRNA concentration [28,34], or inosine-modified-wobble-decoding tRNAs [35] explain the direction of the speed changes in Fig. 1b, we controlled for each of these factors separately by leaving them out of our dataset one at a time, and reapplied Eq. 1 to each new dataset. For example, Green and colleagues identified triplets of amino acids, such as PPG, that slow down translation. To control for these triplets, we removed all amino acid pairs that were found in these specific tripeptide sequence contexts (see SI Methods). In the example of PPG, the reduced data set only has occurrences of the dipeptide PG without an N-terminal flanking proline. We find that even in the absence of these confounding factors, the sign of the speed change remains the same for all 167 pairs and remains the same for 166-out-of-167 pairs when controlling for non-optimal codons (Fig. S3).

To test if wobble base pairing codons versus Watson Crick base pairing codons affect our conclusions, we split our dataset into those instances of amino acid pairs decoded by wobble base pairing and those decoded by Watson Crick geometries. For the wobble-base data set, we find (Fig. S4) that 165-out-of-167 pairs exhibit the same sign change as Fig. 1b. For the Watson Crick dataset 166-out-of-167 pairs exhibit the same sign change as Fig. 1b. Thus, the geometry of the base pairing does not explain our observations in Fig. 1b.

In summary, while these molecular factors undoubtedly contribute to codon translation rates in a variety of contexts, they do not explain the faster or slower amino acid pairs that we observe in Fig. 1b.

### Other robustness tests

There is the potential that other factors associated with our analysis may also influence the results in Fig. 1b. We first tested whether the read-depth threshold (currently requiring at least 3 reads at each codon position) influences our results by constructing data sets with those genes having at least 1 read at (i) 100%, (ii) 90%, and (iii) 75% of the codon positions in their coding sequence, and analyzed these datasets using Eq. 1. We find (Fig. S5) the same sign of the percent change for 166-out-of-167 of the significant amino acids pairs shown Fig. 1b. (Amino acid pair (K,D) switches from slowing down translation to speeding up translation when the read threshold is relaxed to include genes that have less than 75% codons with reads.) Thus, the results presented in Fig. 1b are robust to changes in read-depth threshold.

Next, we tested if mRNA expression level influences our results by splitting the 364 transcripts in our dataset into the half with the highest expression level and the half with the lowest expression level and analyzed each using Eq. 1. We find that the sign change (Fig. S6d, e) is consistent for 166-out-of-167 amino acid pairs indicating that the results in Fig. 1b are not influenced by variation in expression level within our data set.

We then proceeded to test whether broad regions of the transcripts yield different results. To do this, we split our dataset into amino acid pairs, and their associated normalized ribosome densities, from the first half of each coding sequence, the other dataset was from the second half of each coding sequence. Eq. 1 was then applied to each dataset. We find the same sign of the percent change for all 167 significant amino acids pairs in Fig. 1b from the dataset composed of the first half of the coding sequence (Fig. S6b). Similarly, we find the same sign of the percent change for 166-out-of-167 amino acid pairs from the dataset composed of the second half (Fig. S6c). The loss of statistical significance for some of the pairs (lighter orange and lighter blue in Fig. S6) is to be expected due to the reduction of the sample size by half, on average. However, even for these pairs, the sign change is still consistent with Fig. 1b. Thus, our results are not biased by the broad region of the coding sequence from which the ribosome density and amino acid pairs arise from.

Finally, we test whether more localized regions of the coding sequence are biasing our results – including disordered segments, signal peptides, and transmembrane domains. Specifically, we created a dataset in which instances of amino acid pairs arising from disordered protein segments are removed; a dataset in which only cytosolic proteins are present and hence do not contain signal sequences; and a dataset in which regions predicted to be similar to transmembrane domains are removed. Applying Eq. 1 to these datasets we find the sign of the percent change is maintained for all 167 significant amino acid pairs for (Fig. S7). Thus, these local regions of coding sequences are not influencing the results in Fig. 1b.

In summary, the results in Fig. 1b are robust to variation in thresholds used in the analysis, to variation in expression levels, and variation in which regions of the transcript the ribosome profiling reads come from.

### Mutating the P-site amino acid is predicted to alter the translation rate at the A-site

Figure 1b predicts that by keeping the A-site amino acid fixed and mutating the P-site amino acid it is possible to speed up or slow down translation elongation (*i*.*e*., change the sign of the percent change in Eq. 1). For example, when comparing the amino acid pairs (N,R) to (S,R), where R is the amino acid in the A-site, we find (N,R) tends to have 53% more ribosome density than (S,R) (Fig. 1c). Hence, we predict that the codon encoding R in the (S,R) pair will be translated faster than the codon encoding R in the (N,R) pair. We predict that for the 7,980 possible P-site mutations in amino acid pairs where the A-site is fixed there will be a systematic change in translation rate for a majority of them (Fig. 1d and Data S2). Because we are dealing with overlapping ribosome density distributions (Fig. 1c) it is most appropriate to think probabilistically – in terms of the likelihood that a mutation will speed up or slow down translation. We can calculate the odds (Eq. S3) that a mutation at the P-site will speed up or slow down translation. For example, for the mutation (N,R) to (S,R) we calculate translation will speed up with 3-to-1 odds. Thus, out of four randomly selected instances of (N,R) across the proteome, these odds predict that if you mutate N to S for each, three of the instances will speed up translation, and one of the instances will slow down translation on average. We calculated these odds for each of the possible 7,980 mutations and found a broad distribution (Fig. 1e). With odds of 5.7-to-1, mutating (W,G) to (P,G) will slow down translation, while with odds of 1-to-1, mutating (V,W) to (H,W) is as equally likely to speed up translation as it is to slow down translation when Val is mutated to His in the different instances of (V,W) across the proteome (Data S2). In summary, this bioinformatics analysis predicts which P-site amino acid mutations are most likely to result in higher or lower translation elongation rates relative to the wild-type protein sequence.

### Mutational experiments are consistent with amino acid identity influencing translation elongation rates

To experimentally test these predictions, we introduced 12 non-synonymous mutations into various positions of five non-essential *S. cerevisiae* genes that are not involved in translation, and no mutations were made at functional sites of the encoded proteins [36] (Table S2). Five of the mutations are predicted to speed up translation based on Fig. 1b, five are predicted to slow down translation, and two are predicted to have minimal effects on translation speed when the mutated residue is present in the P-site.

To ensure precise measurements at codon resolution we performed ribosome profiling experiments at unconventionally high read depths, having an average of 86 million mapped exome reads per sample after removing reads mapped to rRNA genes, and totaling 1.7 billion mapped exome reads across the samples (Table S3). The resulting ribosome profiles exhibit strong 3-nt periodicity, 87% of mapped reads are in frame zero at a fragment size of 28 nt, and there is a very strong correlation between ribosome profiles for the same gene across samples (Pearson r=0.96, Figs. S8 and S9). These results indicate technical biases are minimal in these experiments, and any such biases that exist will likely cancel out when we carry out the relative comparison between wild type and mutant results.

Comparing the normalized ribosome densities between the wild-type and mutant strains (Fig. 2), we find that in all cases the direction of change in ribosome density at the A-site is consistent with the predictions from Fig. 1b. Two of these ten mutations include a proline in the P-site, for which we observe a speedup when the pair (P,G) is mutated to (E,G) in the gene *YMR122W-A* (Fig. 2a), while mutating (Q,D) to (P,D) in *YOL109W* leads to a slowdown of translation (Fig. 2b). These two mutations serve as positive controls because proline has previously been shown to slow down translation [16,19].

**Figure 2:**
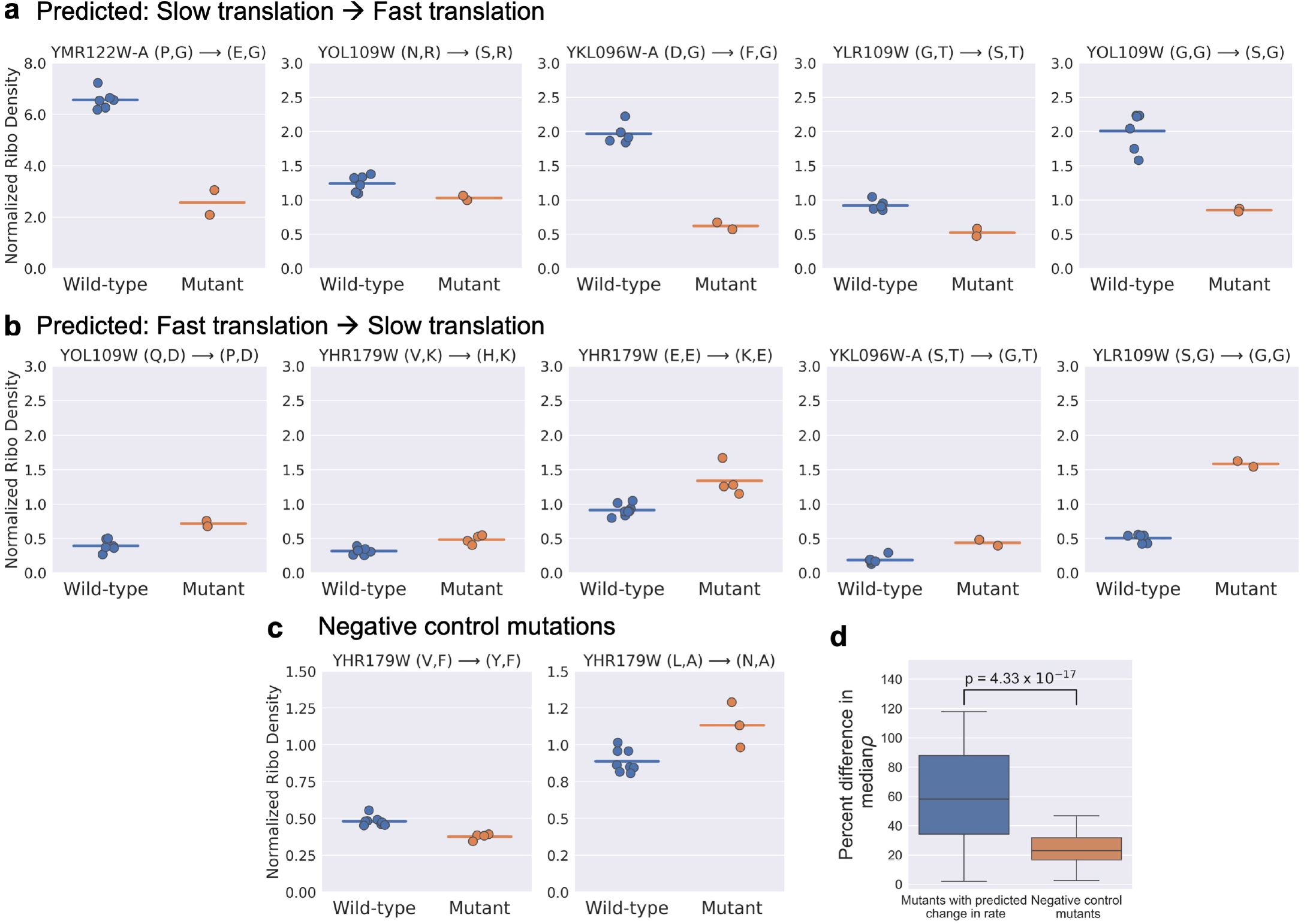
Experiments demonstrate that the identity of amino acids in the P- and A-sites can predictably alter the translation speed of the A-site codon, consistent with the predictions from Fig. 1b. Normalized ribosome density (Eq. S1) upon mutation at five pairs of residues that are predicted to slow down translation **(a)**, five other pairs that are predicted to speed up translation **(b)**, and two negative control mutations that are predicted to have little to no effect on translation speed **(c)** were measured in *S. cerevisiae*. The gene name and pair of amino acids before and after mutation are listed above the panels in (a) through (c). Full details concerning the mutations are provided in Table S2. In each panel, the normalized ribosome density measured at the A-site residue is reported for the wild-type sequence transcript (blue data points) and mutated sequence (orange data points). Each data point corresponds to one biological replicate; the horizontal bar indicates the mean value. The difference between the medians in each panel is statistically significant (one-sided Mann-Whitney U test, *p* = 0.036 for all subpanels in (a) and for mutations in YOL*, YKL* and YLR* in (b). *p* = 0.002 for two mutations in YHR* in (b). *p* = 0.002 and *p* = 0.004 for the two subpanels in (c), respectively). The distribution of percent differences in ribosome density between the mutant and wild-type sequences for the data in panels (a) and (b) is shown as a blue box plot in panel **(d)**, and for the negative control sequences in panel (c), the distribution is shown as an orange box plot in panel (d). The mutations in the negative controls do show a statistically significant difference in normalized densities compared to wild-type (one-sided Mann-Whitney U test, *p* = 0.002 and *p* = 0.004). However, these mutations exhibit a 2.5-fold reduction in effect size (d), consistent with the predictions from Fig. 1b.

Two additional mutations were incorporated as negative controls and are predicted to cause little change in the rate of translation (*i*.*e*., mutations that switch between gray boxes or between the same colored boxes in Fig. 1b). We found that while the normalized ribosome densities of these mutants are statistically different from that of the wild type (Fig. 2c) the effect size is much smaller. The median effect size on translation speed was 2.5-fold lower than what we observed for the other 10 mutations (Fig. 2d). That the negative controls exhibit minor changes in translation speed is to be expected. As previously discussed, we are dealing with overlapping ribosome density and speed distributions (Figs. 1c, e), and there is an associated odds of seeing some speed up or slow down.

In summary, the results from these mutational experiments are consistent with the hypothesis that the P-site amino acid can predictably alter the translation rate at the A-site.

### A qualitative bioinformatics assessment of amino acid versus tRNA contributions

The amino acid mutations we introduced also change the identity of the tRNA molecule at the P-site. Therefore, this change in tRNA identity could be an alternative explanation for the cause of the altered translation speeds (Fig. 2). We have already shown that the decoding geometry is not the main source of the translation rate change, as both Watson-Crick and Wobble base pairing codons yield similar results (Fig. S4). However, it could still be the case that the chemical identity of the tRNA pairs, and the interactions between them, are driving the speed changes. To qualitatively assess whether it is amino acid identity or tRNA identity that is driving the changes we projected the same ribosome profiling data in Fig. 1b onto the 64-by-61 matrix of the codons that can reside in the P- and A-sites. To retain statistical power, we relaxed our gene filtering criteria to include genes that have at least one read at 95% of codon positions in each coding sequence. We then calculated this codon pair matrix for all the 6 datasets and identified the robust codon pairs (Fig. S10a, Data S3) based on the same criteria we used to identify the robust amino acid pairs shown in Fig. 1b (*i*.*e*., the colored boxes). We find that 93% of the robust codon pairs in Fig. S10a have the same sign of the percent change as observed for the robust amino acid pairs they encode in Fig. 1b. Specifically, for each amino acid pair that is significant and robust in Fig. 1b, we asked ‘for those codon pairs that encode a given amino acid pair, and exhibit a statistically significant difference from the average ribosome density, how many of these exhibit the same sign of the speed change as the amino acid pair’? For example, for amino acid pair (D, T), which slows down translation, 7-out-of-8 codon pairs also slow down translation, while the 8^th^ codon pair also slows down translation but is not statistically significant (Fig. S10b). We count this example as being consistent between the amino acid pair and codon pairs, because the loss of statistical power is to be expected when we switch from projecting the data of a 420-element matrix (Fig. 1b) to a 3,904-element matrix (Fig. S10a). Thus, in the vast majority of instances, the amino acid pair translation rate change (Fig. 1b) and codon pair translation rate change (Fig. S10a) are consistent. This indicates that for these instances, it is the amino acid identity primarily driving the speed change because it is consistent across all of the different synonymous codons that are decoded by different tRNA molecules.

For 7% of the codon pairs, however, one or more of the codon pairs exhibit a translation rate change in the opposite direction from the others. For example, for amino acid pair (R,G) in Fig. S10c, the codon pairs with a CGA codon in the P-site leads to slower-than-average translation when GGU and GGA codons are in the A-site, but when CGC and AGG codons are in the P-site there is faster than average translation when GGU is in the A-site. Hence, it seems likely that the interaction of different tRNAs in the P- and A-site for these 12 amino acid pairs primarily drive the translation rate change.

Taken together, the results from this analysis supports the qualitative conclusion that the amino acid pairs often contribute to the sign of the speed change in Fig. 1b, but there are situations where tRNA pairs predominantly drive the speed change.

### A qualitative experimental assessment of amino acid versus tRNA contributions

To experimentally estimate the contribution of amino acid identity versus tRNA identity we took the three mutations that we previously incorporated into the gene *YOL109W* (Table S2) and created a new gene construct with the same three amino acid mutations but that used synonymous codons that are decoded by different tRNA molecules (Table S4). There is a strong correlation between the mutant strains created for this comparison and hence we can compare the normalized ribosome densities at the A-site when these mutations are in the P-site locations of the ribosomes (Fig. S11). For the mutation (N,R) to (S,R), for example, we previously used the codon UCC to mutate N to S. In the new strain, we used the synonymous codon UCG, which is decoded by a different tRNA molecule [37] (Fig. 3a). For the mutants (G,G) to (S,G) and (Q,D) to (P,D), there was a change in normalized ribosome density that was in the same direction and similar in magnitude regardless of the tRNA molecule used (Figs. 3b, c). This indicates that for these mutations, the change in amino acid identity in the P-site is the primary cause of the change in translation rate in the A-site (Fig. 3b, c). In contrast, for the mutation (N,R) to (S,R), we observed a change in ribosome density when one tRNA molecule was used but no change in ribosome density when another tRNA molecule was used (Fig. 3d), indicating that the tRNA identity was the primary cause. Thus, these experimental results support the conclusion that in some cases it is the amino acid identity that causes the change in speed and in others it is the tRNA identity.

**Figure 3:**
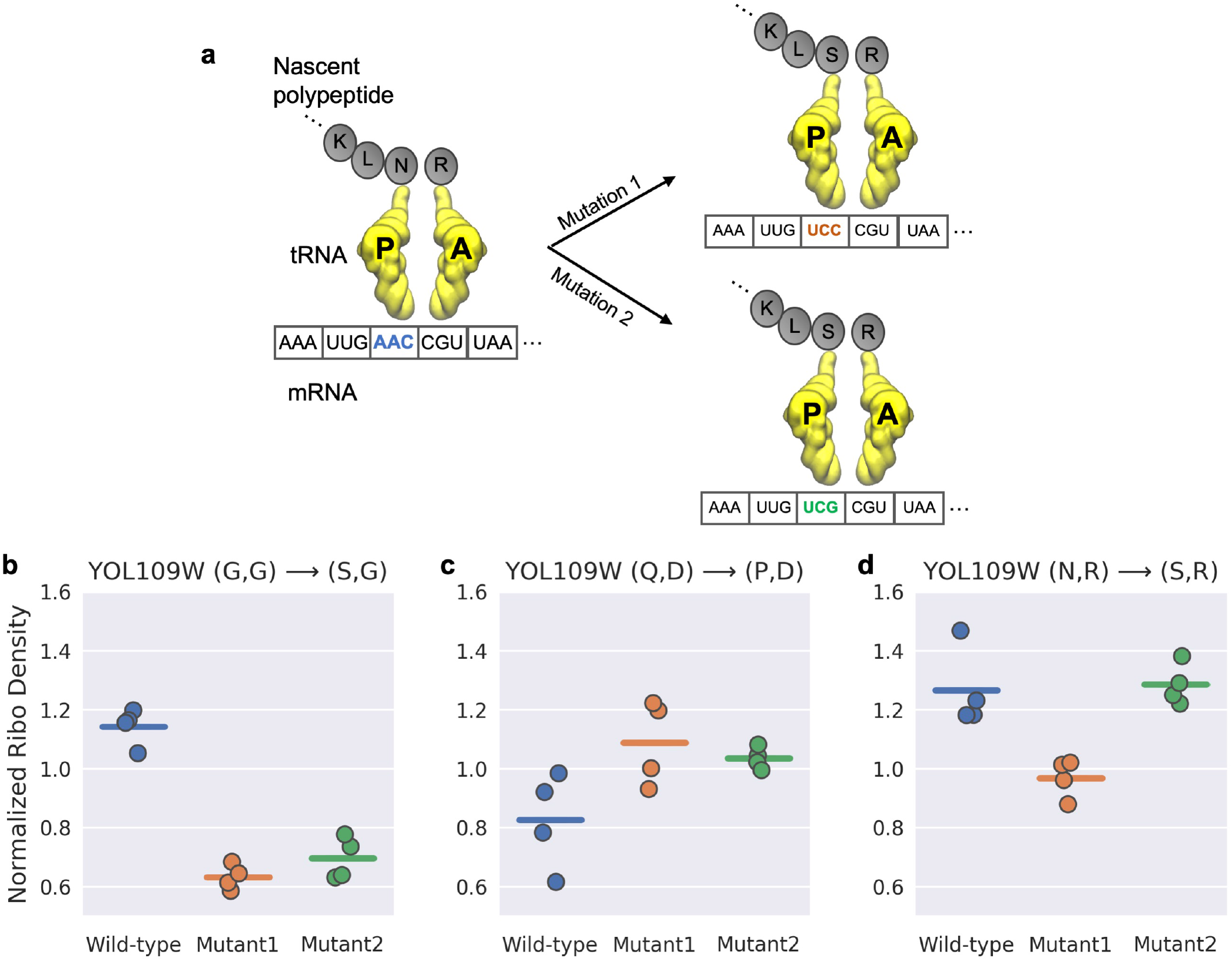
Depending on the amino acid pair, translation speed is influenced by either the identity of the tRNA pair, the amino acid pair, or both. **(a)** For the same amino acid mutation, two mutants are created using synonymous codons decoded by different tRNAs. **(b-d)** To experimentally measure the contribution of amino acid identity, a given non-synonymous mutation in the P-site was encoded using two different synonymous codons, each resulting in the same amino acid mutation but decoded by different tRNA molecules (see (a)). For the mutations *YOL109W* (G,G)→(S,G) (b) and *YOL109W* (Q,D)→(P,D) (c), the change in ribosome density from wild-type was similar for both synonymous mutants (Mutant 1 vs Mutant 2, one-sided Mann-Whitney U test, *p* = 0.1715 and *p* = 0.443, respectively), and hence, the amino acid is the predominant cause for the change in translation speed. For the mutation *YOL109W* (N,R)→ (S,R) (d), the speedup was seen for only one mutant while the other mutant exhibits a normalized ribosome density indistinguishable from that of the wild-type (Wild type *vs*. Mutant 2, one-sided Mann-Whitney U test, *p* = 0.243), indicating in this case that the tRNA identity is the predominant cause for the change in speed up mutation.

### Signatures of evolutionary selection for fast and slow translating amino acid pairs

Translation is an energy-intensive process [38] and the efficiency of translation and protein production are influenced by how quickly ribosomes are released from transcripts. Several studies have suggested that evolution has favored codon optimality in highly expressed genes to enable faster translation and quicker release of ribosomes to increase translation efficiency [8,39,40]. It is possible that evolution can also select for faster translation through mechanisms other than codon optimality. If evolutionary selection pressures have acted to encode translation rate information in the primary structures of proteins through pairs of amino acids, then there should be a non-random distribution of fast and slow-translating pairs of amino acids across the proteome. To test this hypothesis, we calculated the enrichment and depletion of all 400 pairs of amino acids across the *S. cerevisiae* proteome relative to the occurrence expected from a random pairing. We selected the top 20% of the amino acid pairs that were enriched across the proteome and the bottom 20% that were depleted and determined how many of the 86 fast-translating and 81 slow-translating amino acid pairs were present in either of these quintiles (Table S5). The odds ratio of fast-translating pairs being enriched across the proteome and slow-translating pairs being depleted was 4.3 (Eq. S4, *p* = 0.0098, Fisher’s exact test), indicating that selection pressures have indeed selected for the presence of fast-translating pairs and selected against slow-translating pairs (Fig. 4a) across the proteome. This result is consistent with the hypothesis [8,39] that evolution selects for molecular factors that increase the global rate of translation.

**Figure 4:**
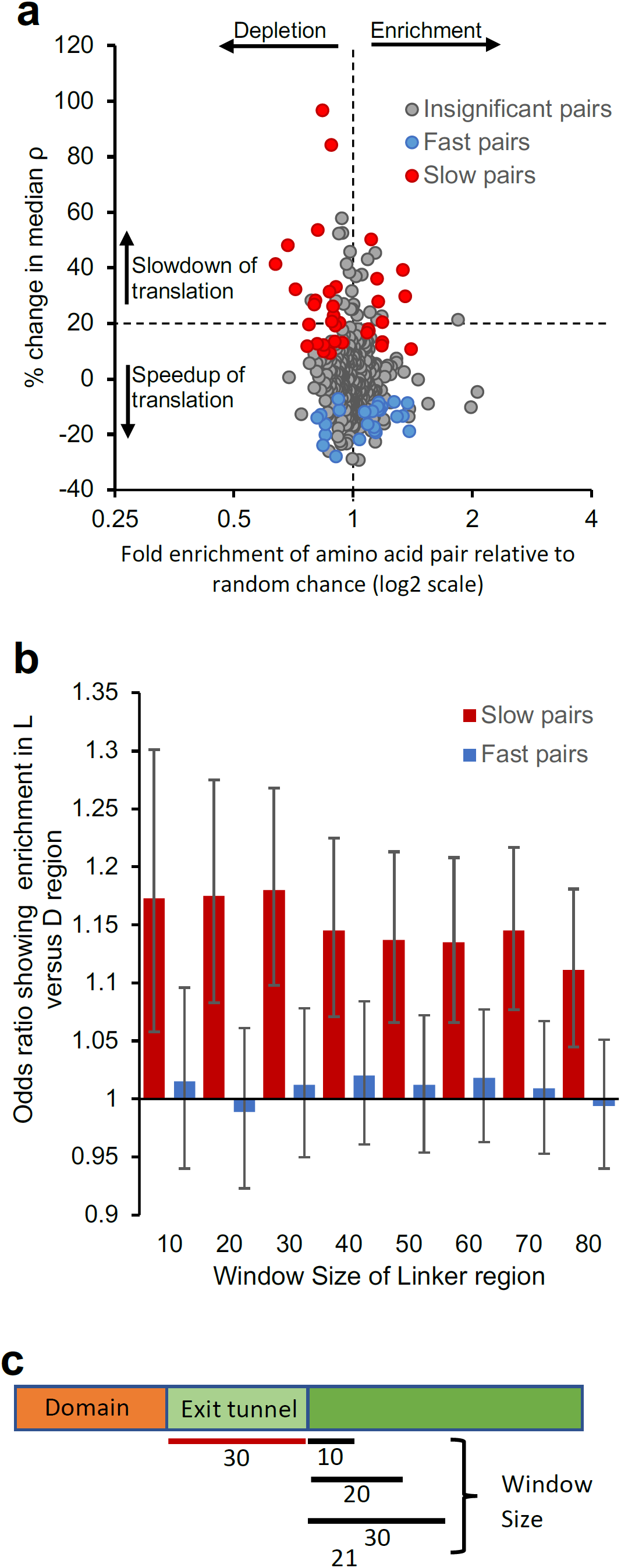
Evolution selects for fast-translating pairs across the proteome but enriches slow-translating pairs across inter-domain linker regions. **(a)** The enrichment and depletion of amino acid pairs across the *S. cerevisiae* proteome is plotted against the percent change in median normalized ribosome densities (ρ) of amino acid pairs taken from Fig. 1b. Among the top 20% enriched and top 20% depleted set, the odds ratio of fast-translating pairs being enriched and slow-translating pairs being depleted is 4.3 (Eq. S4, Fisher’s exact test, *p* = 0.0098). **(b)** The enrichment of fast- and slow-translating pairs in linker (L) regions relative to domain (D) regions. The odds ratio (Eq. S5) gives us a measure of the likelihood of finding either slow or fast pairs in the linker region relative to domain regions compared to the odds of non-slow or non-fast pairs in the linker region relative to domain regions, respectively. An odds ratio greater than 1 would indicate an enrichment in the linker region, while a negative value would indicate a depletion. As a test of robustness, the odds ratio was computed over different window sizes in the linker region, discarding the first 30 residues after the domain to account for those residues being in the ribosome exit tunnel, as illustrated in panel **(c)**. *n* = 170 for a window size of 30 residues. For all window sizes, the odds ratio was significant (Fisher’s exact test, *p* < 0.005 for all linker sizes for slow-translating pairs and insignificant (Fisher’s exact test, *p* > 0.25 for all linker sizes) for fast-translating pairs.

Despite the preference for fast-translating pairs, we found that the slow-translating amino acid pairs were locally enriched by 18% (95% CI: [10%, 27%], *p*=7.9×10^−6^, *n*=170 domains, linker size=30, Fisher’s exact test) in protein segments that are translated after domains have emerged from the ribosome exit tunnel. (Figs. 4b, c). For simplicity, we refer to these segments as linkers. In those linker regions that start 30 residues downstream of domain boundaries, one-fifth of the amino acid pairs, on average, are slow-translating pairs, and in the extreme cases of genes *YDR432W* and *YGL203C*, there are 18 slow pairs in a 30 residue stretch. Codon usage does not explain this enrichment of slow-translating pairs, as we found no difference in the frequency of non-optimal codon usage between linker and domain regions (Fig. S12). These results indicate that a number of slow-translating pairs exist in linker regions that can cumulatively lead to a slowdown of translation as domains fully emerge from the ribosome exit tunnel, which may aid in co-translational folding by providing more time for domains to fold outside of the ribosome exit tunnel.

When the Hsp70 chaperone Ssb is bound to ribosome-nascent chain complexes translation is faster than when Ssb is not bound, possibly because chaperone binding prevents nascent chain folding and hence allows translation to become uncoupled from folding and to proceed faster [41]. We examined if the fast-translating amino acid pairs we identified contributed to this speedup. We found that the fast-translating amino acid pairs were enriched by at least 4% (95% CI: [2.3%, 6.1%], *p*=0.0001, *n*=425, random permutation test) in regions translated while Ssb is bound, suggesting that these pairs do make a contribution (Fig. S13). Taken together, these results indicate that across the primary structures of proteins, evolutionary pressures have selected for amino acid pairs that exhibit faster translation, and along a transcript, fast- and slow-translating pairs are enriched locally in regions that are associated with co-translational folding and chaperone binding.

## DISCUSSION

We have demonstrated that independent of known confounding factors the chemical identity of some pairs of amino acids and tRNA molecules, when present in the P- and A-sites, can predictably and causally result in a speedup or slowdown of translation in the A-site of *S. cerevisiae* ribosomes. An essential and unique feature of our analyses of ribosome profiling data is Eq. 1, which holds fixed the amino acid identity in the A-site while varying the amino acid in the P-site. This approach controls for variation in tRNA concentration and accommodation rates in the A-site, and measures the relative difference in ribosome density when a particular amino acid pair is present in the A- and P-sites compared to when any other amino acid is in the P-site while holding the A-site constant. A recent study [42] estimated the effect of codon and amino acid identity in the P-site but they did not find any effect since they do not control for the other confounding factors that can influence translation rate at the A-site. Hence, our approach (Eq. 1) allows us to isolate the effect of the P-site amino acid on the overall translation rate at the A-site. In total, we identified 167 amino acid pairs that exhibit above average or below average translation speeds (Fig. 1b) and verified these predictions experimentally for 10 of these pairs. These speed changes are not explained by alternative hypotheses concerning the presence of mRNA structure, wobble base pairing, tripeptide motifs, positively charged upstream nascent chain residues, variation in cognate tRNA concentration, differences in expression level, nor other aspects of the analysis (Figs. S3-S4). These speed changes are also robust to changes in thresholds for gene selection criteria as well as choosing instances of amino acid pairs from different regions of the gene transcripts (Fig. S5-S7). As an important, naturally occurring internal control, these results are consistent with the commonly seen translational slow-down effect of proline [16,32] evidenced by the vertical stripe of red-shifted colors in Fig. 1b.

Changes in elongation rate can influence protein structure, function, and cellular phenotype without altering protein expression levels. A variety of studies [4,6,43–50] have demonstrated for different proteins that the total amount of soluble protein produced in a cell, and the total amount of soluble, *functional* protein produced can be very different. The influence of elongation rate on protein structure and function arises in part on the impact translation speed changes have on co-translational processes including domain folding [8]. Thus, the translation elongation speed differences we observe between pairs of amino acids are more likely to influence co-translational processes than expression level.

Over- or under-representation of particular pairs of codons, compared to random chance, occurs across the transcriptome of many organisms in a phenomenon referred to as “codon pair utilization bias” [51]. Some codon pairs occur less frequently as they can result in negative functional consequences. For example, codon pairs with the pattern of nnUAnn [52] can cause a frameshift resulting in a premature stop codon. Hence, evolution has selected against codon pairs that have this motif. The biological importance of codon pair bias is also highlighted in bioengineering applications where de-optimization of codon pairs across the coding sequences of viruses has been used to attenuate their virulence [53]. A biochemical study found that synonymously changing codon pairs results in a change in protein expression level – indicating these pairs are influencing elongation speed [54]. This, and other studies [55–57], hypothesized that synonymous codon pairs translate at different speeds due to differences in the molecular interactions between neighboring tRNAs when they are co-located at the A- and P-sites of the ribosome. The results of this study find that indeed, a change in translation speed upon a change in codon pair in some cases is due to a change in the identity of the tRNA at the P-site, but also that there are many other cases where the change of translation speed is due to the change in identity of the amino-acid at the P-site (Fig. 3b, c). Thus, the results from this study indicate that there is a greater richness of molecular mechanisms that can contribute to codon pair utilization bias and identify a large number of amino-acid pairs where systematic speed differences are observed.

Most studies on the relationship between translation kinetics and nascent peptide sequence features at the A- and P-site have involved the presence of proline, often in the context of ‘*X*PP*Z*’ peptide motifs [15,58,59], where *X* and *Z* can be any amino acid. To observe an effect of these motifs on translation, engineered bacterial cells must be used that lack elongation-factor P because its presence eliminates proline-induced stalling [17,18,60]. *In vitro* enzymology studies involving proline have focused only on the kinetics of the tRNA accommodation and peptide bond formation sub-steps of the translation cycle [16,32]. In contrast, this study used wild-type *S. cerevisiae* cells in which we discovered predictable changes in speed in 138 pairs of amino acids that do not involve proline. The ribosome profiling signal we analyze is proportional to the dwell time of the ribosome at the A-site [23,61], and thus the effects we observe reflect what happens to the overall rate of one cycle of translation elongation rather than just the effect on individual steps of accommodation, peptide bond formation, or translocation. Thus, in contrast to an *in vitro* enzymology study that concluded that although the P-site amino acid can alter the peptide bond formation rate it has little effect on elongation due to the slower tRNA accommodation step [62], we observe wide spread P-site identity effects on overall codon translation rates *in vivo*. This suggests the possibility that for some of these pairs accommodation is not always rate limiting *in vivo* and that the effects of peptide bond formation rates may constitute a larger proportion of the dwell of a ribosome than previously thought. One promising approach to measure this proportion comes from the use of three different antibiotics that can trap *S. cerevisiae* ribosomes in different states of elongation [63]. The depth of coverage in the datasets from that study, with 7 million mapped reads on average compared to 87 million mapped reads in this study, are too sparse for us to test this hypothesis.

We observe consistent and robust influences of the amino-acid at the P-site. The effect of the P-site amino acid on translation speed tends to be consistent in either speeding up/slowing down translation, or having no effect at all, regardless of the amino acid in the A-site. Out the twenty amino acids that can reside in the P-site, nine (A,E,F,I,M,S,T,V,Y) speed up translation (green-shifted colors in Fig. 1b) or have little effect (gray colors); six (D,G,H,K,N,P) slow down translation (red-shifted colors) or have little effect; and three (L,R,W) can either speed up, slow down, or have no effect relative to the average translation speed. For example, Ala in the P-site speeds up translation when A, G, P or Q is in the A-site, Lys in the P-site slows down translation when A, D, E, or Q is in the A-site, whereas Leu in the P-site slows down translation when G or Q is in the A-site but speeds up translation when D, K or N is in the A-site. Thus, 15-out-of-20 amino acids, when present in the P-site, consistently change translation speed in one direction. Pro, Glu, Ser, Gly and Asp are the most robust in terms of changing speed for the largest number of possible A-site amino acids, as evidenced by their vertical stripes of colors in Fig. 1b. Consistent with our findings, an *in vitro* enzymology study [62] that measured peptide bond formation rates of puromycin with eight different types of amino acids in the P-site found that Asp was the second slowest amino acid to form a peptide bond with puromycin after proline, and serine was the second fastest.

We find that when the P-site amino acid is mutated in many cases it is the amino acid identity change that drives the translation speed change, but we also find instances where it is the tRNA change that is the driver. Pairs whose effect arise primarily due to tRNA identity could influence steps in the translation-elongation cycle other than peptide bond formation, including hybrid state formation and translocation as these steps structurally alter the intermolecular interactions between tRNA molecules, and between tRNA molecules and the ribosome. Disentangling the detailed molecular causes of the speed changes identified in Fig. 1b is likely to be a fruitful area of future research. Furthermore, in our analyses we averaged out the effect of the tRNAs that reside in the E-site, as Eq. 1 is only conditioned on what is in the A- and P-sites. Two kinetic studies [64,65] demonstrated that a small number of tRNA’s in the E-site can slow mRNA translocation through the ribosome and contribute to frameshifting and stop codon read-through. Thus, exploring the influence of the E-site tRNA on translation at A-site using ribosome profiling data is an exciting area for future research.

Our evolutionary analyses indicate a proteome-wide selection for fast-translating amino acid pairs, potentially to increase the efficiency of energy-intensive process of translation [66] while locally enriching slow-translation pairs that might aid in co-translational processes important for a protein to attain its structural and functional form. Evolutionary selection pressures select only against phenotypic traits, not genotype. Therefore, the enrichment of fast-translating amino acid pairs across the *S. cerevisiae* transcriptome and the clusters of slow- and fast-translating pairs along transcripts that are correlated with co-translational processes suggest that the elongation kinetics encoded by these pairs influence organismal phenotype and fitness. More speculatively, these results open up the possibility that there may exist disease-causing amino acid mutations that do not alter the final folded structures of proteins but instead alter the co-translational behavior and processing of the nascent proteins via altered elongation kinetics.

In summary, separate from other molecular factors known to influence translation speed, elongation kinetics are causally and predictably encoded in protein primary structures through the identity of particular pairs of amino acids and the tRNAs they are attached to, with broad implications for protein and mRNA sequence evolution, and translational control of gene expression.

## ACKNOWLEDGMENTS

Computations for this research were performed on the Pennsylvania State University’s Institute for Computational and Data Sciences’ Advanced CyberInfrastructure (ICS-ACI). This work was supported by a DKFZ NCT3.0 Integrative Project in Cancer Research (NCT3.0_2015.54 DysregPT), a European Research Council Advanced grant (TransFold 743118), and the Deutsche Forschungsgemeinschaft (SFB 1036) to B.B, a National Institutes of Health MIRA R35 (Project number 1R35GM124818-01) and National Science Foundation ABI (Innovation Award 1759860) to E.P.O.

## AUTHOR CONTRIBUTIONS

E.P.O. conceived the study. U.F., G.K. and B.B. carried out experiments to generate *S. cerevisiae* mutant strains and Ribosome profiling. N.A. analyzed the data. N.A., P.S., P.C. contributed to analysis methods of published ribosome profiling data and annotation of domain boundaries in *S. cerevisiae*. N.S.A contributed to all statistical analyses and their interpretation. N.A, P.S., U.F., G.K. and E.P.O. wrote the manuscript.

## DECLARATION OF INTERESTS

The authors declare no competing interests.

## DATA AVAILABILITY

Mutant ribosome profiling sequencing data is available on NCBI GEO database with accession number GSE133370.

The source code to generate the results and key figures has been made available through a Jupyter notebook that can be accessed at https://github.com/obrien-lab/P-site-A-site-pairs-analysis/

## Notes

### Competing Interest Statement

The authors have declared no competing interest.

## REFERENCES

[1] P. Shah, Y. Ding, M. Niemczyk, G. Kudla, J.B. Plotkin, Rate-limiting steps in yeast protein translation., Cell. 153 (2013) 1589–601. https://doi.org/10.1016/j.cell.2013.05.049.

[2] A. Espah Borujeni, H.M. Salis, Translation Initiation is Controlled by RNA Folding Kinetics via a Ribosome Drafting Mechanism, J. Am. Chem. Soc. 138 (2016) 7016–7023. https://doi.org/10.1021/jacs.6b01453.

[3] D.A. Nissley, E.P. O’Brien, Timing Is Everything: Unifying Codon Translation Rates and Nascent Proteome Behavior, J. Amercan Chem. Soc. 136 (2014) 17892–17898. https://doi.org/10.1021/ja510082j.

[4] M. Zhou, J. Guo, J. Cha, M. Chae, S. Chen, J.M. Barral, M.S. Sachs, Y. Liu, Non-optimal codon usage affects expression, structure and function of clock protein FRQ, Nature. 495 (2013) 111–115. http://dx.doi.org/10.1038/nature11833.

[5] I.M. Sander, J.L. Chaney, P.L. Clark, Expanding Anfinsen’s Principle : Contributions of Synonymous Codon Selection to Rational Protein Design Expanding Anfinsen’s Principle : Contributions of Synonymous Codon Selection to Rational Protein Design, J. Am. Chem. Soc. 136 (2014) 858–861.

[6] G. Zhang, M. Hubalewska, Z. Ignatova, Transient ribosomal attenuation coordinates protein synthesis and co-translational folding., Nat. Struct. Mol. Biol. 16 (2009) 274–280. https://doi.org/10.1038/nsmb.1554.

[7] A.K. Sharma, E.P. O’Brien, Non-equilibrium coupling of protein structure and function to translation–elongation kinetics, Curr. Opin. Struct. Biol. 49 (2018) 94–103. https://doi.org/10.1016/J.SBI.2018.01.005.

[8] K.C. Stein, J. Frydman, The stop-and-go traffic regulating protein biogenesis: How translation kinetics controls proteostasis, J. Biol. Chem. 294 (2018) 2076–2084. https://doi.org/10.1074/jbc.rev118.002814.

[9] J.L. Chaney, P.L. Clark, Roles for Synonymous Codon Usage in Protein Biogenesis, Annu. Rev. Biophys. 44 (2015) 143–166. https://doi.org/10.1146/annurev-biophys-060414-034333.

[10] T. Ikemura, Codon usage and tRNA content in unicellular and multicellular organisms., Mol. Biol. Evol. 2 (1985) 13–34. https://doi.org/10.1093/oxfordjournals.molbev.a040335.

[11] J. Choi, R. Grosely, A. Prabhakar, C.P. Lapointe, J. Wang, J.D. Puglisi, How Messenger RNA and Nascent Chain Sequences Regulate Translation Elongation, Annu. Rev. Biochem. 87 (2018) 421–449. https://doi.org/10.1146/annurev-biochem-060815-014818.

[12] X. Qu, J.-D. Wen, L. Lancaster, H.F. Noller, C. Bustamante, I. Tinoco, The ribosome uses two active mechanisms to unwind messenger RNA during translation., Nature. 475 (2011) 118–121. https://doi.org/10.1038/nature10126.

[13] T. Tuller, Y.Y. Waldman, M. Kupiec, E. Ruppin, Translation efficiency is determined by both codon bias and folding energy, Proc. Natl. Acad. Sci. 107 (2010) 3645–3650. https://doi.org/10.1073/pnas.0909910107.

[14] A.K. Sharma, P. Sormanni, N. Ahmed, P. Ciryam, U.A. Friedrich, G. Kramer, E.P. O’Brien, A chemical kinetic basis for measuring translation initiation and elongation rates from ribosome profiling data, PLOS Comput. Biol. 15 (2019) e1007070. https://doi.org/10.1371/journal.pcbi.1007070.

[15] A.P. Schuller, C.C.C. Wu, T.E. Dever, A.R. Buskirk, R. Green, eIF5A Functions Globally in Translation Elongation and Termination, Mol. Cell. 66 (2017) 194–205.e5. https://doi.org/10.1016/j.molcel.2017.03.003.

[16] M.Y. Pavlov, R.E. Watts, Z. Tan, V.W. Cornish, M. Ehrenberg, A.C. Forster, Slow peptide bond formation by proline and other N-alkylamino acids in translation., Proc. Natl. Acad. Sci. U. S. A. 106 (2009) 50–54. https://doi.org/10.1073/pnas.0809211106.

[17] S. Ude, J. Lassak, A.L. Starosta, T. Kraxenberger, D.N. Wilson, K. Jung, Translation elongation factor EF-P alleviates ribosome stalling at Polyproline Stretches, Science. 339 (2013) 82–85. https://doi.org/10.1126/science.1228985.

[18] L.K. Doerfel, I. Wohlgemuth, C. Kothe, F. Peske, H. Urlaub, M. V. Rodnina, EF-P Is Essential for Rapid Synthesis of Proteins Containing Consecutive Proline Residues, Science. 339 (2013) 85–88. https://doi.org/10.1126/science.1229017.

[19] C.G. Artieri, H.B. Fraser, Accounting for biases in riboprofiling data indicates a major role for proline in stalling translation, Genome Res. 24 (2014) 2011–2021. https://doi.org/10.1101/gr.175893.114.

[20] C.A. Charneski, L.D. Hurst, Positively Charged Residues Are the Major Determinants of Ribosomal Velocity, PLoS Biol. 11 (2013) e1001508. https://doi.org/10.1371/journal.pbio.1001508.

[21] R.D. Requião, H.J.A. de Souza, S. Rossetto, T. Domitrovic, F.L. Palhano, Increased ribosome density associated to positively charged residues is evident in ribosome profiling experiments performed in the absence of translation inhibitors, RNA Biol. 13 (2016) 561–568. https://doi.org/10.1080/15476286.2016.1172755.

[22] J. Lu, C. Deutsch, Electrostatics in the Ribosomal Tunnel Modulate Chain Elongation Rates, J. Mol. Biol. 384 (2008) 73–86. https://doi.org/10.1016/j.jmb.2008.08.089.

[23] N.T. Ingolia, S. Ghaemmaghami, J.R.S. Newman, J.S. Weissman, Genome-wide analysis in vivo of translation with nucleotide resolution using ribosome profiling., Science. 324 (2009) 218–223. https://doi.org/10.1126/science.1168978.

[24] N.T. Ingolia, Ribosome Footprint Profiling of Translation throughout the Genome, Cell. 165 (2016) 22–33. https://doi.org/10.1016/j.cell.2016.02.066.

[25] D.A. Nissley, A.K. Sharma, N. Ahmed, U.A. Friedrich, G. Kramer, B. Bukau, E.P. O’Brien, Accurate prediction of cellular co-translational folding indicates proteins can switch from post-to co-translational folding, Nat. Commun. 7 (2016) 10341. https://doi.org/10.1038/ncomms10341.

[26] C.C. Williams, C.H. Jan, J.S. Weissman, Targeting and plasticity of mitochondrial proteins revealed by proximity-specific ribosome profiling, Science. 346 (2014) 748–751. https://doi.org/10.1126/science.1257522.

[27] C.H. Jan, C.C. Williams, J.S. Weissman, “Principles of ER cotranslational translocation revealed by proximity-specific ribosome profiling,” Science. 346 (2014) 748–751. https://doi.org/10.1126/science.1257521.

[28] D.E. Weinberg, P. Shah, S.W. Eichhorn, J.A. Hussmann, J.B. Plotkin, D.P. Bartel, Improved Ribosome-Footprint and mRNA Measurements Provide Insights into Dynamics and Regulation of Yeast Translation, Cell Rep. 14 (2016) 1787–1799. https://doi.org/10.1016/j.celrep.2016.01.043.

[29] D.J. Young, N.R. Guydosh, F. Zhang, A.G. Hinnebusch, R. Green, Rli1/ABCE1 Recycles Terminating Ribosomes and Controls Translation Reinitiation in 3′UTRs In Vivo, Cell. 162 (2015) 872–884. https://doi.org/10.1016/j.cell.2015.07.041.

[30] J.A. Hussmann, S. Patchett, A. Johnson, S. Sawyer, W.H. Press, Understanding Biases in Ribosome Profiling Experiments Reveals Signatures of Translation Dynamics in Yeast, PLoS Genet. 11 (2015) e1005732. https://doi.org/10.1371/journal.pgen.1005732.

[31] A. Diament, T. Tuller, Estimation of ribosome profiling performance and reproducibility at various levels of resolution, Biol. Direct. 11 (2016) 24. https://doi.org/10.1186/s13062-016-0127-4.

[32] M. Johansson, K.-W. Ieong, S. Trobro, P. Strazewski, J. Aqvist, M.Y. Pavlov, M. Ehrenberg, pH-sensitivity of the ribosomal peptidyl transfer reaction dependent on the identity of the A-site aminoacyl-tRNA, Proc. Natl. Acad. Sci. 108 (2010) 79–84. https://doi.org/10.1073/pnas.1012612107.

[33] J. Der Wen, L. Lancaster, C. Hodges, A.C. Zeri, S.H. Yoshimura, H.F. Noller, C. Bustamante, I. Tinoco, Following translation by single ribosomes one codon at a time, Nature. 452 (2008) 598–603. https://doi.org/10.1038/nature06716.

[34] A. Dana, T. Tuller, The effect of tRNA levels on decoding times of mRNA codons, Nucleic Acids Res. 42 (2014) 9171–9181. https://doi.org/10.1093/nar/gku646.

[35] C.E. Gamble, C.E. Brule, K.M. Dean, S. Fields, E.J. Grayhack, Adjacent Codons Act in Concert to Modulate Translation Efficiency in Yeast, Cell. 166 (2016) 679–690. https://doi.org/10.1016/j.cell.2016.05.070.

[36] J. Cherry, E. Hong, C. Amundsen, R. Balakrishnan, G. Binkley, C. ET, C. KR, C. MC, D. SS, E. SR, F. DG, H. JE, H. BC, K. K, K. CJ, M. SR, N. RS, P. J, S. MS, S. M., W. S, W. ED, Saccharomyces Genome Database: the genomics resource of budding yeast., Nucleic Acids Res. 40 (2012) D700–5.

[37] J. Hani, H. Feldmann, tRNA genes and retroelements in the yeast genome, Nucleic Acids Res. 26 (1998) 689–696. https://doi.org/10.1093/nar/26.3.689.

[38] D.M. Stoebel, A.M. Dean, D.E. Dykhuizen, The cost of expression of Escherichia coli lac operon proteins is in the process, not in the products, Genetics. 178 (2008) 1653–1660. https://doi.org/10.1534/genetics.107.085399.

[39] C.E. Brule, E.J. Grayhack, Synonymous Codons: Choose Wisely for Expression, Trends Genet. 33 (2017) 283–297.

[40] A.A. Komar, The Yin and Yang of codon usage., Hum. Mol. Genet. 25 (2016) R77–R85. https://doi.org/10.1093/hmg/ddw207.

[41] K. Döring, N. Ahmed, T. Riemer, H.G. Suresh, Y. Vainshtein, M. Habich, J. Riemer, M.P. Mayer, E.P. O’Brien, G. Kramer, B. Bukau, Profiling Ssb-Nascent Chain Interactions Reveals Principles of Hsp70-Assisted Folding, Cell. 170 (2017) 298–311.e20. https://doi.org/10.1016/j.cell.2017.06.038.

[42] R. Tunney, N.J. McGlincy, M.E. Graham, N. Naddaf, L. Pachter, L.F. Lareau, Accurate design of translational output by a neural network model of ribosome distribution, Nat. Struct. Mol. Biol. 25 (2018) 577–582. https://doi.org/10.1038/s41594-018-0080-2.

[43] A. a. Komar, A pause for thought along the co-translational folding pathway, Trends Biochem. Sci. 34 (2009) 16–24. https://doi.org/10.1016/j.tibs.2008.10.002.

[44] F. Buhr, S. Jha, M. Thommen, J. Mittelstaet, F. Kutz, H. Schwalbe, M. V Rodnina, A.A. Komar, Synonymous Codons Direct Cotranslational Folding toward Different Protein Conformations., Mol. Cell. 61 (2016) 341–351. https://doi.org/10.1016/j.molcel.2016.01.008.

[45] W. Holtkamp, G. Kokic, M. Jäger, J. Mittelstaet, A.A. Komar, M. V Rodnina, Cotranslational protein folding on the ribosome monitored in real time., Science. 350 (2015) 1104–1107. https://doi.org/10.1126/science.aad0344.

[46] M. V Rodnina, The ribosome in action: Tuning of translational efficiency and protein folding., Protein Sci. 25 (2016) 1390–1406. https://doi.org/10.1002/pro.2950.

[47] P.S. Spencer, E. Siller, J.F. Anderson, J.M. Barral, Silent substitutions predictably alter translation elongation rates and protein folding efficiencies., J. Mol. Biol. 422 (2012) 328–335. https://doi.org/10.1016/j.jmb.2012.06.010.

[48] C.-J. Tsai, Z.E. Sauna, C. Kimchi-Sarfaty, S. V Ambudkar, M.M. Gottesman, R. Nussinov, Synonymous mutations and ribosome stalling can lead to altered folding pathways and distinct minima., J. Mol. Biol. 383 (2008) 281–291. https://doi.org/10.1016/j.jmb.2008.08.012.

[49] S.J. Kim, J.S. Yoon, H. Shishido, Z. Yang, L.A. Rooney, J.M. Barral, W.R. Skach, Protein folding. Translational tuning optimizes nascent protein folding in cells., Science. 348 (2015) 444–448. https://doi.org/10.1126/science.aaa3974.

[50] I. Fedyunin, L. Lehnhardt, N. Böhmer, P. Kaufmann, G. Zhang, Z. Ignatova, tRNA concentration fine tunes protein solubility, FEBS Lett. 586 (2012) 3336–3340. https://doi.org/10.1016/j.febslet.2012.07.012.

[51] G.A. Gutman, G.W. Hatfield, Nonrandom utilization of codon pairs in Escherichia coli, Proc. Natl. Acad. Sci. 86 (1989) 3699–3703. https://doi.org/10.1073/pnas.86.10.3699.

[52] A. Tats, T. Tenson, M. Remm, Preferred and avoided codon pairs in three domains of life., BMC Genomics. 9 (2008) 463. https://doi.org/10.1186/1471-2164-9-463.

[53] J.R. Coleman, D. Papamichail, S. Skiena, B. Futcher, E. Wimmer, S. Mueller, Virus Attenuation by Genome-Scale Changes in Codon Pair Bias, Science (80-.). 320 (2008) 1784 LP – 1787. https://doi.org/10.1126/science.1155761.

[54] B. Irwin, J.D. Heck, G.W. Hatfield, Codon pair utilization biases influence translational elongation step times, J. Biol. Chem. 270 (1995) 22801—22806. https://doi.org/10.1074/jbc.270.39.22801.

[55] J.R. Buchan, L.S. Aucott, I. Stansfield, tRNA properties help shape codon pair preferences in open reading frames, Nucleic Acids Res. 34 (2006) 1015–1027. https://doi.org/10.1093/nar/gkj488.

[56] D. Smith, M. Yarus, tRNA-tRNA interactions within cellular ribosomes, Proc. Natl. Acad. Sci. U. S. A. 86 (1989) 4397–4401. https://doi.org/10.1073/pnas.86.12.4397.

[57] K.H. Nierhaus, J. Wadzack, N. Burkhardt, R. Jünemann, W. Meerwinck, R. Willumeit, H.B. Stuhrmann, Structure of the elongating ribosome: Arrangement of the two tRNAs before and after translocation, Proc. Natl. Acad. Sci. U. S. A. 95 (1998) 945–950. https://doi.org/10.1073/pnas.95.3.945.

[58] L. Peil, A.L. Starosta, J. Lassak, G.C. Atkinson, K. Virumae, M. Spitzer, T. Tenson, K. Jung, J. Remme, D.N. Wilson, Distinct XPPX sequence motifs induce ribosome stalling, which is rescued by the translation elongation factor EF-P, Proc. Natl. Acad. Sci. 110 (2013) 15265–15270. https://doi.org/10.1073/pnas.1310642110.

[59] A.L. Starosta, J. Lassak, L. Peil, G.C. Atkinson, K. Virumäe, T. Tenson, J. Remme, K. Jung, D.N. Wilson, Translational stalling at polyproline stretches is modulated by the sequence context upstream of the stall site, Nucleic Acids Res. 42 (2014) 10711–10719. https://doi.org/10.1093/nar/gku768.

[60] C.J. Woolstenhulme, N.R. Guydosh, R. Green, A.R. Buskirk, High-Precision analysis of translational pausing by ribosome profiling in bacteria lacking EFP, Cell Rep. 11 (2015) 13–21. https://doi.org/10.1016/j.celrep.2015.03.014.

[61] N.T. Ingolia, G.A. Brar, S. Rouskin, A.M. McGeachy, J.S. Weissman, The ribosome profiling strategy for monitoring translation in vivo by deep sequencing of ribosome-protected mRNA fragments, Nat. Protoc. 7 (2012) 1534–1550. https://doi.org/10.1038/nprot.2012.086.

[62] I. Wohlgemuth, S. Brenner, M. Beringer, M. V. Rodnina, Modulation of the rate of peptidyl transfer on the ribosome by the nature of substrates, J. Biol. Chem. 283 (2008) 32229–32235. https://doi.org/10.1074/jbc.M805316200.

[63] C.C.C. Wu, B. Zinshteyn, K.A. Wehner, R. Green, High-Resolution Ribosome Profiling Defines Discrete Ribosome Elongation States and Translational Regulation during Cellular Stress, Mol. Cell. 73 (2019) 959–970.e5. https://doi.org/10.1016/j.molcel.2018.12.009.

[64] J. Choi, J.D. Puglisi, Three tRNAs on the ribosome slow translation elongation., Proc. Natl. Acad. Sci. U. S. A. 114 (2017) 13691–13696. https://doi.org/10.1073/pnas.1719592115.

[65] C. Chen, B. Stevens, J. Kaur, Z. Smilansky, B.S. Cooperman, Y.E. Goldman, Allosteric vs. spontaneous exit-site (E-site) tRNA dissociation early in protein synthesis., Proc. Natl. Acad. Sci. U. S. A. 108 (2011) 16980–16985. https://doi.org/10.1073/pnas.1106999108.

[66] J.B. Plotkin, G. Kudla, Synonymous but not the same: the causes and consequences of codon bias., Nat. Rev. Genet. 12 (2011) 32–42. https://doi.org/10.1038/nrg2899.

