## Supplementary material for "Pairs of amino acids at the P- and A-sites of the ribosome predictably and causally modulate translation-elongation rates": SI

**Materials and Methods:**

**Details of Experiments**

***Design of mutant strains***

There are 7,980 possible amino acid mutations for all combinations of amino acid pairs where the P-site amino acid can be mutated while keeping the A-site amino acid unchanged. Bioinformatic analyses of published ribosome profiling data predicted that 4,134 out of these 7,980 possible mutations will result in a significant change in speed (Fig. 1d and Data S2). To experimentally validate our bioinformatic predictions, we chose 5 mutations that are predicted to accelerate translation and 5 mutations that are predicted to retard translation. Two more mutations were created where our bioinformatic analysis predicted no significant change in speed. These 12 mutations were chosen such that they represent as many different combinations of amino acids and also in such a manner that they can be mutated on a small number of genes. Mutations (P,G) → (E,G) and (Q,D) → (P,D) were chosen to act as positive control since Proline has been known to slow down translation when present in P-site. (G,G) → (S,G) and (S,G) → (G,G) were implemented to test the complementarity of the mutations, i.e., if mutating the P-site from G → S is having a significant change in translation speed, is S → G having the same effect in the opposite direction? To experimentally verify whether the effect on speed for G → S and S → G are also possible for more than one A-site, we carry our similar mutations with T in the A-site, i.e., the mutations (G,T) → (S,T) and (S,T) → (G,T). The rest of the mutations were chosen to represent amino acids not represented in the above mutations. Location of these mutations were chosen such that the normalized ribosome density in the published datasets at these instances of the amino acid pair is close to the median and an instance is avoided that is at the extreme tail of the distribution. The mutations were chosen on 5 non-essential highly expressed genes where these mutations can be distributed on the instances of these amino acid pairs. The chosen genes were not involved in the process of ribosome biogenesis or translation. The mutated positions on the selected genes were chosen in order that they were not at the functional sites or sites subjected to post-translational modifications as defined in the Saccharomyces Genome Database [1]. The gene name and location of mutations are listed in Table S2.

We denote the five mutant strains as YKL*, YMR*, YLR*, YOL* and YHR* that were created so that each strain contained mutations in a single gene: *YKL096W-A*, *YMR122W-A*, *YLR109W*, *YOL109W* and *YHR179W* respectively. To assess the effect of tRNA versus amino acid identity on translation, an additional mutant strain, denoted YOL**, was created which contained the same amino acid mutations as YOL*, but using a synonymous set of codons. Details concerning these two set of synonymous mutations is provided in Table S4.

Ribosome profiling was carried out in two phases. In the first phase, two replicates of mutant strains YKL*, YMR*, YLR* and YOL* were subject to ribosome profiling. For the single mutation in *YKL096W-A*, the mutant ribosome densities were from two replicates of YKL* while the wild-type ribosome densities were from YMR*, YLR* and YOL* which contained the endogenous transcript for *YKL096W-A*. A similar procedure was followed for the four other mutations where we predict a speedup of translation and three mutations where we predicted a slowdown of translation, present on these four genes.

In the second phase, ribosome profiling was run for four replicates of YHR*, YOL* and YOL**. YOL* and YOL** contained the same amino acid mutations but differed in terms of the set of synonymous codons used. YHR* contained four mutations. Two mutations were the negative control mutations. The other two were mutations predicted to retard translation bringing the total mutations where we predict a slowdown of translation to 5. The 8 samples (4 replicates each of YOL* and YOL**) were used as wild type for mutations in *YHR179W* while the 4 replicates of YHR* served as wild-type samples containing the endogenous *YOL109W* gene against the *YOL109W* mutations in YOL* and YOL**. The number of replicates was increased to 4 in the second phase to generate enough sample size for a valid statistical test. The normalized ribosome densities were not compared across the two phases as the ribosome profiling samples prepared on different days generally show poor correlation of ribosome densities at the codon level (Fig. S14).

#### *Strain Construction of mutants*

A two-step procedure omitting selection markers in the final construct was used for mutant strain construction. First, the gene of interest was replaced in the strain BY4741 by a *K. lactis URA3* cassette according to Janke *et al.* [2]. Second, the desired mutant gene enclosing overhangs (45nt/60nt) was constructed by PCR and used to replace the introduced *URA3* cassette by homologous recombination. Candidates were selected on 5-Fluoroorotic Acid (5-fluorouracil-6-carboxylic acid monohydrate; 5-FOA) containing plates and insertion of the correct mutations was verified by colony PCR and DNA sequencing.

#### *Ribosome profiling and library preparation*

200 mL of cells were grown in YPD to an $OD_{600\ nm}$ of 0.5, rapidly filtered (All-Glass Filter 90mm, Millipore), flash frozen in liquid nitrogen, mixed with 600 µL frozen lysis buffer (20 mM Tris-HCl pH 8.0, 140 mM KCl, 6 mM MgCl2, 0.1% NP-40, 0.1 mg/ml CHX, 1 mM PMSF, 2x Complete EDTA-free protease inhibitors (5056489001, Roche), 0.02 U/ml DNase I (4716728001, Roche), 20 mg/mL leupeptin, 20 mg/mL aprotinin, 10 mg/mL E-64, 40 mg/mL bestatin) and pulverized by mixer milling (2 min, 30 Hz, MM400, Retsch). Thawed cell lysates were cleared by centrifugation (20,000xg, 5 min, 4°C) and digested by RNase I (AM2295, Ambion; 125 U/1mg nucleic acid) for 1 hr (25°C, 650 rpm) to obtain ribosome footprints. The reaction was stopped by adding 10 µl SUPERase-In (AM2696, Ambion). The digested lysate was loaded onto 10-50% (w/v) sucrose gradients and centrifuged at 35,000 rpm, 4°C for 2.5 hrs. Gradients were fractionated and monosome fractions were collected, pooled and used for RNA purification by hot acid-phenol extraction. 5 µg of purified RNA was depleted for rRNA using the Ribo-Zero Gold for Yeast kit (MRZY1306, Illumina). Deep sequencing libraries were prepared following the protocol described in Döring *et al.* [3] and sequenced on a HiSeq 2000 (Illumina).

### Computational analyses of ribosome profiling data

#### *Analysis of ribosome profiling datasets*

Wild-type ribosome profiling datasets were obtained from five different published studies [4–8] whose accession numbers are provided in Table S1. The raw reads for each of these published datasets were preprocessed according to the steps specified in the Methods of the respective study. The sequenced reads for all ribosome profiling datasets of mutant strains prepared for this study were subject to a uniform preprocessing step. The raw reads were first trimmed of their 3′

adapter sequence CTGTAGGCACCATCAATTCGTATGCCGTCTTCTGCTTG using cutadapt v1.14 [9]. The reads were also subject to a quality filter of at least 20 during the cutadapt run.

For all downstream analyses, a uniform protocol was used as specified below. Preprocessed reads were first mapped to a set of ribosomal RNA sequences using Bowtie2 [10] and then subsequently the unmapped reads were mapped to the rest of *S. cerevisiae* reference genome sacCer3 using Tophat2 [11]. Custom python scripts were implemented for all downstream analyses. Mapped reads were first quantified by their 5′ ends on individual gene transcripts and A-site positions were assigned according to Table 1 in Suppl. Ref. [12]. To maintain the accuracy of read assignment, transcripts in which multiple mapped reads constitute more than 0.1% of the reads mapped to the CDS region were not considered in the analysis. To minimize noise and increase the confidence in the results, we restrict our dataset to only transcripts that have at least 3 reads mapped at every codon position for the dataset with the highest coverage (Williams' dataset). Applying this filter, we obtain 364 genes for Williams' dataset [4], which has the highest coverage among the published datasets. Hence, we use this dataset for all downstream analyses.

***Estimation of translation speed change for amino acid pairs***

The normalized ribosome density $\rho$ for every codon position $j$ in transcript $i$ is calculated by dividing the number of mapped ribosome profiling reads $R_{k,i}$ by the average reads mapped to the transcript $i$ consisting of $N_{C,i}$ codons.

$$\rho_{j,i} = \frac{R_{j,i}}{\sum_k R_{k,i}/N_{C,i}} \quad \text{[S1]}$$

$\rho$ values are binned into an individual distribution for every amino acid pair $(X,Z)$ where $X$ is in the P-site and $Z$ is in the A-site. The distribution [$\rho(X,Z)$] is populated by $\rho(j,i)$ for each codon position $j$ in transcript $i$ such that $(j-1_{AA}, j_{AA}) = (X,Z)$. The terms $j$ and $i$ are dropped from [$\rho(X,Z)$] since this is an aggregated distribution of all instances of $\rho_{j,i}$ for the amino acid pair $(X,Z)$. The speedup or slowdown of translation caused by an amino acid pair is estimated using the percent change in median of the distribution [$\rho(X,Z)$] as compared to median of the distribution [$\rho(\sim X,Z)$] described in Eq [1] and restated below

$$Percent\ change\ = \frac{Median[\rho(X,Z)] - Median[\rho(\sim X,Z)]}{Median[\rho(\sim X,Z)]} * 100\ \%. \quad \text{[1]}$$

$(\sim X,Z)$ represents the set of all pairs of amino acids where $Z$ is in the A-site and $X$ is not in the P-site. A positive percent change (red shades in Fig. 1b) will indicate that for amino acid pair $(X,Z)$, presence of $X$ is leading to slower translation (higher values of $\rho$) of $Z$ as compared to when $X$ is not present in the P-site. A negative percent change (green shades in Fig. 1b) will indicate $Z$ is translated faster when $X$ is present in the P-site as compared to when $X$ is no present in the P-site.

The distribution [$\rho(X,Z)$] is plotted in Fig. 1c for two pairs (N,R) and (S,R). $\rho$ is plotted across the X-axis and the probability density $P(\rho)$ is plotted on the Y-axis that is calculated below as

$$P(\rho(X,Z)) = \frac{\sum_i \sum_{j=2} \rho_{(j-1,j)\,=\,(X,Z)}\left(\Theta(\rho_{j,i}-\delta_\rho)-\Theta(\rho_{j,i}+\delta_\rho)\right)}{2\delta_\rho \sum_i \sum_{k=2} \rho_{(k-1,k)\,=\,(X,Z)}} \quad \text{[S2]}$$

where $\delta_\rho$ is half the bin width used to construct the histogram. $\Theta(\rho_{j,i} - \delta_\rho)$ and $\Theta(\rho_{j,i} + \delta_\rho)$ are terms of Heaviside step function that are used to classify whether the term $\rho_{j,i}$ is to be included in a particular bin of width $\delta_\rho$ or not. The factor of $2\delta_\rho$ is the bin-width of the histogram. By dividing by $2\delta_\rho$ we make $P(\rho(X,Z))$ a probability density function.

An odds measure is calculated for whether a P-site mutation $(X,Z)$ → $(B,Z)$ or $(B,Z)$ → $(X,Z)$ will result in a change of translation rate. If we assume that $(X,Z)$ is a slow-translating pair and $(B,Z)$ is a fast-translating pair, then we calculate the odds of how likely it is for any instance of $(X,Z)$ when mutated to $(B,Z)$ will lead to speedup of translation. For the odds calculation, we first compute a difference matrix of dimension MxN, where M is the number of instances of $(X,Z)$ in our dataset, and N is the number of instances of $(B,Z)$. Each element of the lower triangle of the matrix is the difference in ribosome density measured for a specific instance of $(X,Z)$ minus a specific instance of $(B,Z)$. The ratio of positive differences to negative differences of these values (Eq. S3) reports the odds that a speedup will occur when $(X,Z)$ is mutated to $(B,Z)$. This odds is equivalent to odds of a slowdown when any instance of $(B,Z)$ is mutated to $(X,Z)$.

$$Odds = \frac{N\{\rho_{(X,Z)} - \rho_{(B,Z)}\}>0}{N\{\rho_{(X,Z)} - \rho_{(B,Z)}\}<0}, \qquad Median(\rho_{(X,Z)}) > Median\,(\rho_{(B,Z)}) \qquad [S3]$$

#### *Statistical significance, correction for false discoveries, and reproducibility of the trends across different datasets*

Normalized ribosome density profiles were calculated from the six ribosome profiling data sets for the 364 high coverage genes identified in the Williams dataset [4]. $[\rho(X,Z)]$ and $[\rho(\sim X,Z)]$ were then calculated for all pairs of amino acids in the P- and A-sites. Instances of zero A-site reads within these datasets were not included in the distributions. The Mann-Whitney U test is used to assess statistical significance between these normalized ribosome density distributions of amino acid pairs. In the 20X21 matrix (shown in Fig. S2 and Fig. 1b), we are carrying out 420 pair comparisons and hence to control for false discovery, Benjamini Hochberg FDR correction is applied to the 420 uncorrected p-values resulting in 420 corrected p-values that determine the statistical significance of the pairs which are shown as colored boxes in Fig. S2 for each of the six datasets. The percent change in median normalized ribosome density $\rho$ is considered to be reproducible if it is positive (slowdown) or negative (speedup) in all 6 datasets and statistically significant in at least 4 out of the 6 datasets. These robust and significant pairs are shown as colored boxes in Fig. 1b. This criterion accounts for the false discovery proportion across these datasets and overcomes any deficiencies of Benjamini-Hochberg false discovery rate correction.

#### *Controlling for molecular factors that could influence the results in Fig. 1b*

In Figs. S3-S7, we test whether the effects observed in Fig. 1b arise from molecular factors other than the amino acid identity of the pair. To test if a potentially confounding factor affects our conclusions, we remove from our dataset all the instances in which that confounding factor is present, apply Eq. 1 to the resulting dataset, visualize the results as in Fig. 1b, and determine if there was a change in the sign of the percent change in the list of robust amino acid pairs we identified in Fig. 1b. If there was not a change, then we conclude that factor is not influencing our conclusions.

Below, we list the selection criteria for the molecular factors and how they are applied:

**(1)** The *in vivo* mRNA secondary structure information was obtained from DMS-Seq dataset for *S. cerevisiae* [13] and instances with structured nucleotides present 4 to 6 codons downstream of A-site were removed.

**(2)** For the effect of upstream positively charged residues, we identified those amino acid pairs that had 1 or more positive charged within 2 to 5 residues upstream of the amino acid pair in the A- and P-site and removed them from our dataset.

**(3)** To test if codon optimality is having an effect, we remove instances of amino acid pairs encoded by non-optimal codons at A-site. The list of non-optimal codons we used is defined by Pechmann and Frydman 2013 [14]

**(4)** To test for the effect of proline-containing and other stalling tripeptide motifs, we remove instances of amino acid pairs that are present in the context of these tripeptide motifs. For example, PPG is one of the stalling motif and if we are considering the effect of PG and if the amino acid upstream of the P-site is a P leading to peptide sequence of PPG, we remove this instance of PG from the distribution. The list of stalling motifs are obtained from Suppl. Ref. [15] and defined as those having pause scores in wild-type samples as greater than 3.

**(5)** To test for Watson Crick versus Wobble base pairing, we divide our distribution into two subsets. One whose instances of amino acid pairs are encoded by codons through Watson-Crick base pairing only and the other whose instances of amino acid pairs are encoded by codons through Wobble base pairing. The list of codons that get decoded through Watson-Crick base pairing and Wobble base pairing are reported in Table 2 of Suppl. Ref. [16] where Watson-Crick codons have a cognate tRNA while Wobble codons do not have a cognate tRNA.

**(6)** To test for expression level, we rank ordered, from highest to lowest, the 364 genes in the dataset based on expression (RPKM mapped to entire transcript) and divided the list into two equal sized subsets of 182 genes – the top half are highly expressed, the bottom half are low expressed. Eq. 1 was then applied to each dataset separately.

**(7)** The disordered protein regions were obtained from Table S7 of Suppl. Ref. [17]. Instances of amino acid pairs from these regions were removed from our dataset and Eq. 1 applied to the remaining data.

**(8)** To test if properties of membrane proteins like presence of signal peptides or trans membrane domains may be driving our effect, we carry out our analysis by restricting ourselves to transcripts that encode only cytosolic proteins. The localization information is obtained from Sacchromyces Genome Database (SGD) [1]. Regions with properties of trans membrane domains are also predicted across non-membrane proteins. We identified them from SGD and remove any instances from these regions for this control analysis.

#### *Statistical significance for the mutational experiments*

One-sided Mann-Whitney U test is applied to determine the statistical significance of the difference between the normalized ribosome densities of mutant and wild type strains in Figs. 2 and 3. For the five mutations in Fig. 2a and three mutations in Fig. 2b, $n = 2$ and $m$ =6 for mutant and wild-type respectively and the distributions of their normalized ribosome densities are distinct with no overlapping data points giving us the smallest p-value of 0.036 for comparison of distributions with the small sample sizes. For the 2 mutations in YHR179W in Fig. 2b (in gene YHR179W), $n = 4$ for mutants and $m = 8$ for wild-types and the distributions are also distinct giving us the smallest p-value of 0.002. In Fig. 3, pairwise comparisons are made between the three sets of Wild-type, Mutant1 and Mutant 2 whose respective p-values are listed in the caption of Fig. 3.

#### *Enrichment/Depletion of amino acid pairs*

Enrichment/Depletion of an amino acid pair across the proteome of *S. cerevisiae* is calculated by dividing the observed probability of finding the amino acid pair by the probability expected of forming these pairs by random chance, which is the product of the probabilities of the individual amino acids across the proteome. This ratio is a measure of enrichment/depletion of the amino acid pair which we call the enrichment score. To test for evolutionary selection of amino acid pairs that significantly influence translation speed, the top 20% (80 out of 400 amino acid pairs) of amino acid pairs are taken that have the highest enrichment scores (highly enriched across the transcriptome) and the bottom 20% of amino acid pairs with the lowest enrichment scores (highly depleted across the transcriptome). Fisher's exact test is used to test the hypothesis that the fast pairs are more likely to be enriched while slow pairs are more likely to be depleted across the *S. cerevisiae* transcriptome. The odds ratio of fast pairs being enriched and slow pairs being depleted was calculated as:

$$Odds\ ratio = \frac{Enriched(Fast\ amino\ acid\ pairs) \times Depleted(Slow\ amino\ acid\ pairs)}{Enriched\ (\ Slow\ amino\ acid\ pairs) \times Depleted(Fast\ amino\ acid\ pairs)} \qquad \text{[S4]}$$

#### *Classification of downstream linker and domain regions*

A database of 864 *S. cerevisiae* proteins with annotated domain boundaries was created using the same procedure as in Suppl. Ref. [18]. This database is available in Data S4. Domain regions are defined as existing between the starting and ending codon positions reported in Data S4. Inter-domain linker regions are defined as those regions between domains. And for the analysis presented in Fig. 4, the enrichment analysis was carried out using the domain regions, and only those linker regions starting 30 residues downstream of the C-terminal domain boundary. This accounts for the approximately 30 linker residues needed to span the exit tunnel and sterically permit tertiary domain folding outside the exit tunnel. If the linker is shorter than the sum of the 30 residues in the tunnel and a window size (Fig. 4c), then this domain-linker pair is not considered in the analysis for this particular window size. The enrichment/depletion of fast and slow amino acid pairs is calculated in the linker region relative to the domain region in terms of odds ratio and the statistical significance is estimated using Fisher's exact test. Significant pairs with stop codon in A-site are not considered in this analysis. The instances for the remaining 78 slow-translating pairs and 85 fast-translating pairs determined within the defined linker and domain regions. The enrichment is calculated for the slow-translating amino acid pairs by odds ratio measure as shown below in Eq. S5. Similarly, odds ratio can be calculated for fast-translating amino acid pairs compared to non-fast translating amino acid pairs.

$$Odds\ ratio = \frac{N(Slow\ amino\ acid\ pairs\ in\ linker\ ) \times N(Non-slow\ amino\ acid\ pairs\ in\ domain)}{N\ (\ Slow\ amino\ acid\ pairs\ in\ domain) \times N(Non-slow\ amino\ acid\ pairs\ in\ linker)} \quad [S5]$$

***Enrichment/depletion of amino acid pairs in Ssb-bound translated regions***

In a previous study [3], regions of mRNA transcripts were identified that are translated by the ribosome when Hsp70 chaperone Ssb is bound to the nascent polypeptide chain and when it is not bound. A Fold Enrichment (FE) measure is the ratio of selective ribosome profiling reads to ribosome profiling reads and its profile across an mRNA transcript, where FE is above a threshold, defines the regions that are translated when Ssb is bound. We define 5 thresholds based on the percentile values of FEs from the Cumulative Distribution Function of FE values (see Fig S6 from study of Döring *et al.* [3]). For example, for thresholds ($P_{80}$, $P_{20}$), every nucleotide position with a FE value greater than $P_{80}$ was classified as Ssb-bound translated region (B region) while every position with a FE value lower than $P_{20}$ was defined as Ssb-unbound translated region (UB region). The rest of the nucleotides with FE value between $P_{20}$ and $P_{80}$ were ignored. Similarly, 5 thresholds are defined to represent the increasing differential of Ssb binding strength as we move from left to right on X-axis in Fig. S13.

In the study of Döring *et al.* [3], it was shown that translation was faster, on average, in the Ssb-bound translated regions relative to Ssb-unbound regions. The presence or absence of several molecular factors, such as downstream mRNA secondary structure, optimal codons, and proline content were shown to correlate with the increased translation speed in the Ssb-bound regions. To test whether amino acid pairs are also contributing as a molecular factor, we test the hypothesis that fast amino acid pairs are enriched and slow amino acid pairs are depleted across Ssb-bound translated regions relative to Ssb-unbound translated regions. Permutation test is used to measure the statistical significance of the percent change of probabilities of fast and slow amino acid pairs in Ssb-bound translated region (B region) relative to Ssb-unbound translated regions (UB region) which is plotted on Y-axis is Fig. S13. Error bars are calculated using bootstrapping [19].

a

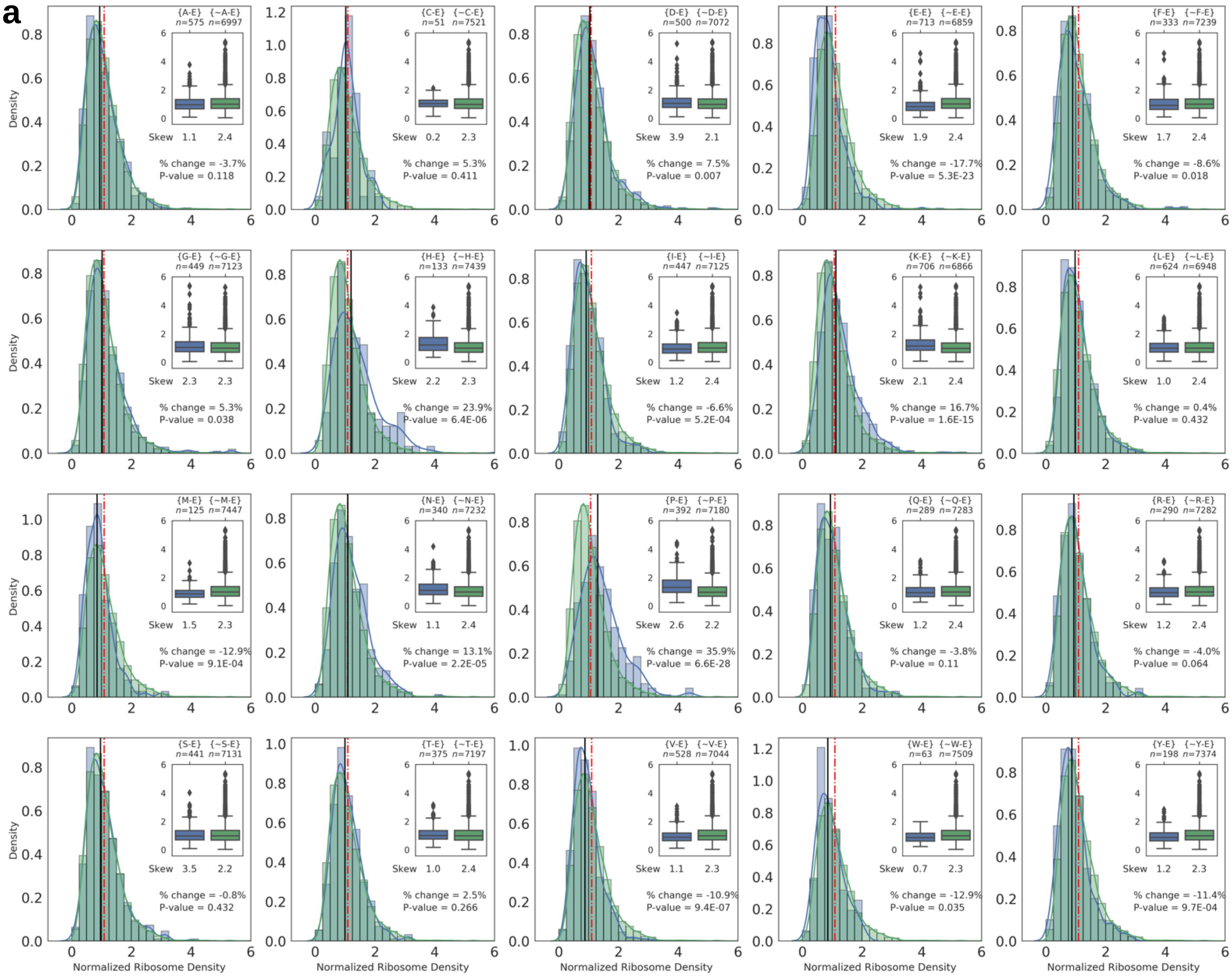
{A-E} {~A-E}
n=575 n=6997
Skew 1.1 2.4
% change = -3.7%
P-value = 0.118
{C-E} {~C-E}
n=51 n=7521
Skew 0.2 2.3
% change = 5.3%
P-value = 0.411
{D-E} {~D-E}
n=500 n=7072
Skew 3.9 2.1
% change = 7.5%
P-value = 0.007
{E-E} {~E-E}
n=713 n=6859
Skew 1.9 2.4
% change = -17.7%
P-value = 5.3E-23
{F-E} {~F-E}
n=333 n=7239
Skew 1.7 2.4
% change = -8.6%
P-value = 0.018
{G-E} {~G-E}
n=449 n=7123
Skew 2.3 2.3
% change = 5.3%
P-value = 0.038
{H-E} {~H-E}
n=133 n=7439
Skew 2.2 2.3
% change = 23.9%
P-value = 6.4E-06
{I-E} {~I-E}
n=447 n=7125
Skew 1.2 2.4
% change = -6.6%
P-value = 5.2E-04
{K-E} {~K-E}
n=706 n=6866
Skew 2.1 2.4
% change = 16.7%
P-value = 1.6E-15
{L-E} {~L-E}
n=624 n=6948
Skew 1.0 2.4
% change = 0.4%
P-value = 0.432
{M-E} {~M-E}
n=125 n=7447
Skew 1.5 2.3
% change = -12.9%
P-value = 9.1E-04
{N-E} {~N-E}
n=340 n=7232
Skew 1.1 2.4
% change = 13.1%
P-value = 2.2E-05
{P-E} {~P-E}
n=392 n=7180
Skew 2.6 2.2
% change = 35.9%
P-value = 6.6E-28
{Q-E} {~Q-E}
n=289 n=7283
Skew 1.2 2.4
% change = -3.8%
P-value = 0.11
{R-E} {~R-E}
n=290 n=7282
Skew 1.2 2.4
% change = -4.0%
P-value = 0.064
{S-E} {~S-E}
n=441 n=7131
Skew 3.5 2.2
% change = -0.8%
P-value = 0.432
{T-E} {~T-E}
n=375 n=7197
Skew 1.0 2.4
% change = 2.5%
P-value = 0.266
{V-E} {~V-E}
n=528 n=7044
Skew 1.1 2.3
% change = -10.9%
P-value = 9.4E-07
{W-E} {~W-E}
n=63 n=7509
Skew 0.7 2.3
% change = -12.9%
P-value = 0.035
{Y-E} {~Y-E}
n=198 n=7374
Skew 1.2 2.3
% change = -11.4%
P-value = 9.7E-04
Density
Normalized Ribosome Density

**b**

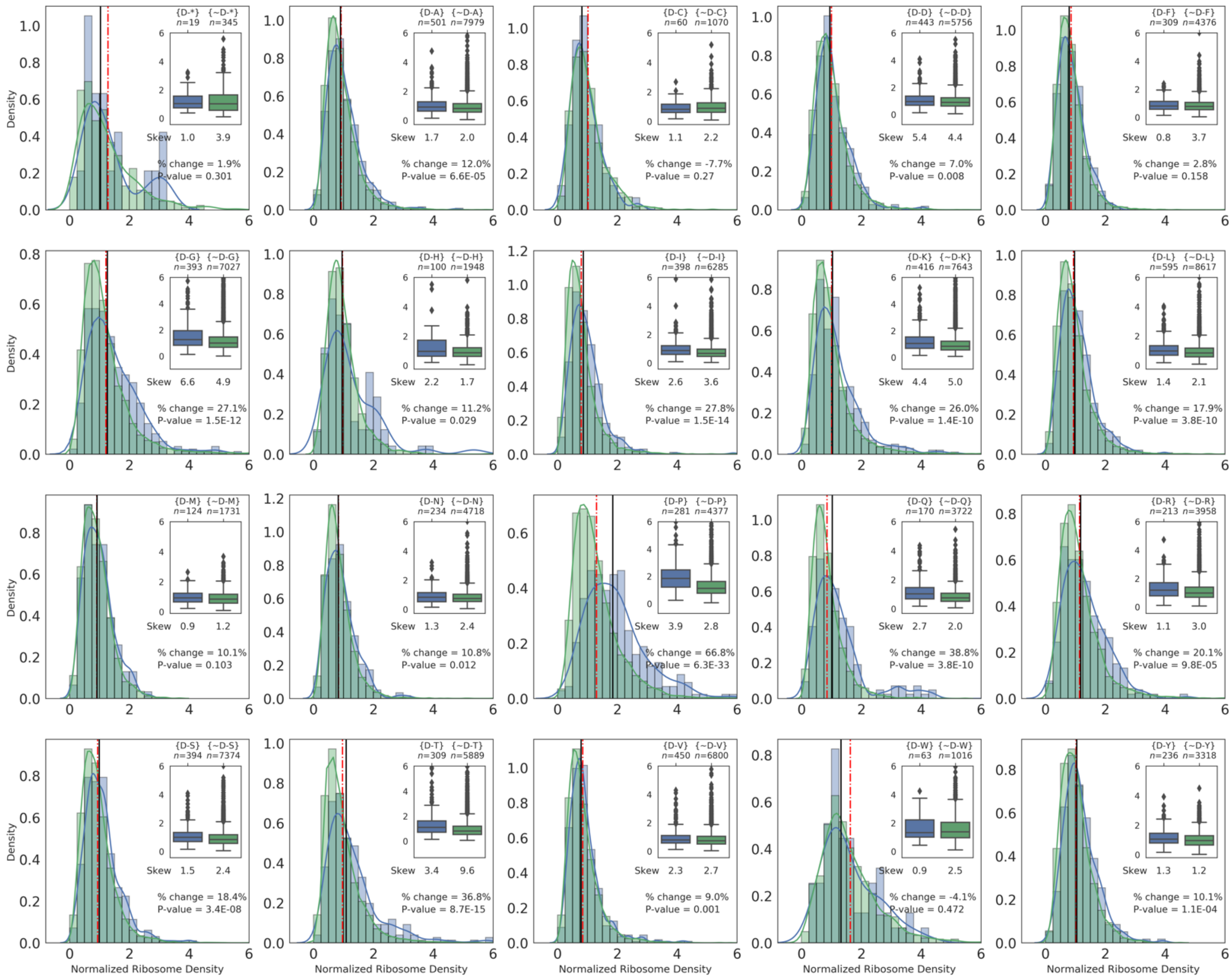


**Figure S1: Normalized ribosome density distributions (Eq. S2) for amino acid pair $(X, Z)$ and $(\sim X, Z)$ for the Williams dataset. (a)** For amino acid E in the A-site, there are 20 amino acids that it can be paired with in the P-site. These 20 distributions are plotted for $(X, E)$ and $(\sim X, E)$. The median values are shown as vertical lines $(X, E)$ (black solid line) and $(\sim X, E)$ (red dotted line) while the percent change in translation rate (Eq. 1) and the corresponding p-value along with the skew of the distributions are listed. For visualization purposes, the range of normalized ribosome density has been limited to a value of 6. These results constitute the row for A-site amino acid E in the matrix presented in Fig. S2a **(b)** 21 distributions are possible for amino acid P in the P-site with 20 amino acid in the A-site and also stop codon in the A-site. Plot for $(D, E)$ has been shown in Fig. S1a. The other 20 possible distributions are plotted here whose results constitute the column for P-site amino acid D in the matrix presented in Fig. S2a. The content of each plot is same as (a).

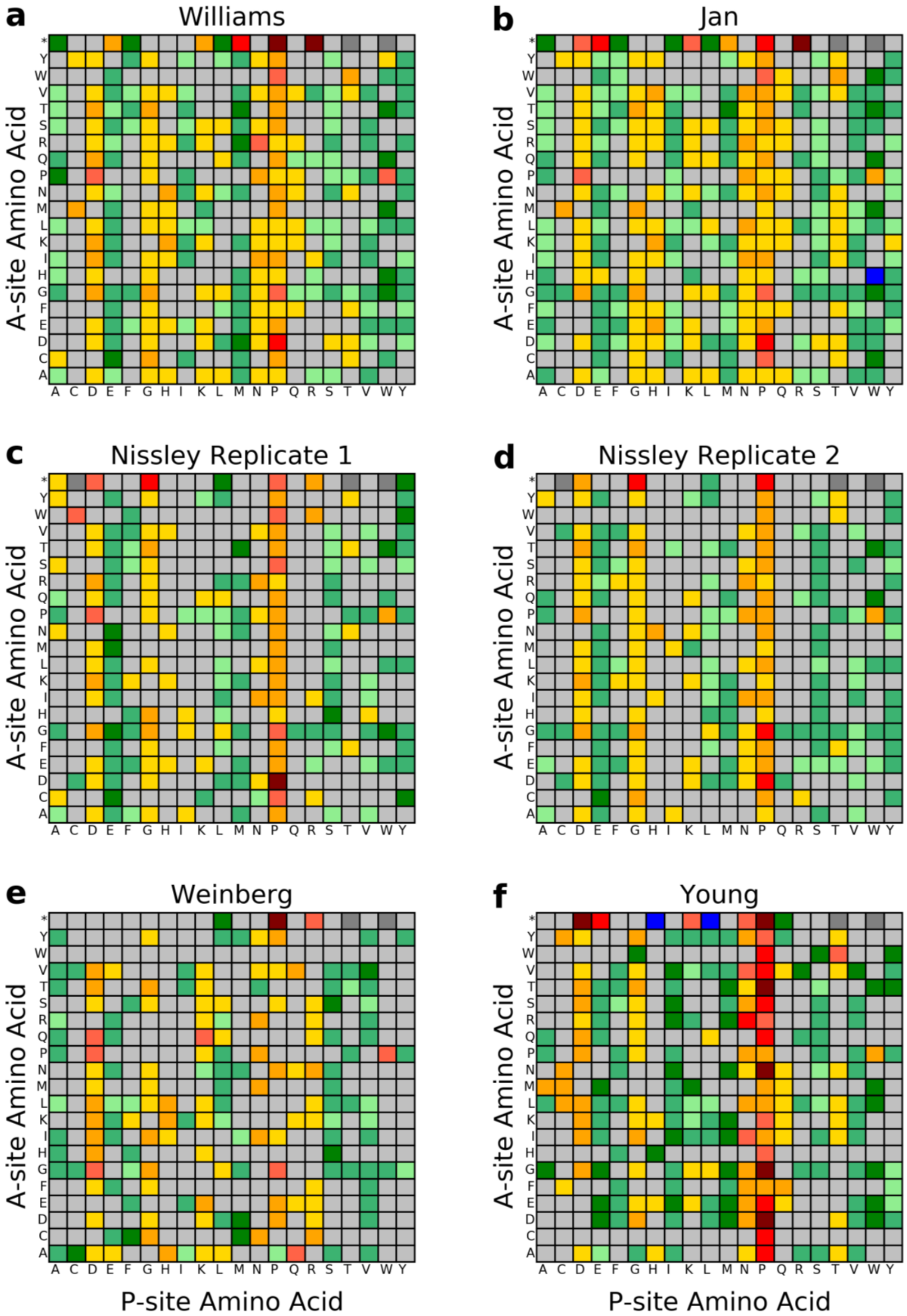
a
Williams
b
Jan
c
Nissley Replicate 1
d
Nissley Replicate 2
e
Weinberg
f
Young
A-site Amino Acid
P-site Amino Acid
* Y W V T S R Q P N M L K I H G F E D C A
A C D E F G H I K L M N P Q R S T V W Y

**Figure S2: The percent change in median normalized ribosome density ρ for a given pair of amino acids in the P-site and A-site, relative to any other amino acid being in the P-site (Eq. 1).** The published datasets are listed based on the name of the first author of the study, namely, Williams [4] **(a)**, Jan [6] **(b)**, two biological replicates from study of Nissley [5] **(c, d)**, Weinberg [7] **(e)** and Young [8] **(f)**. The accession numbers of these samples are listed in Table S1. The legend is the same as in Fig. 1b.

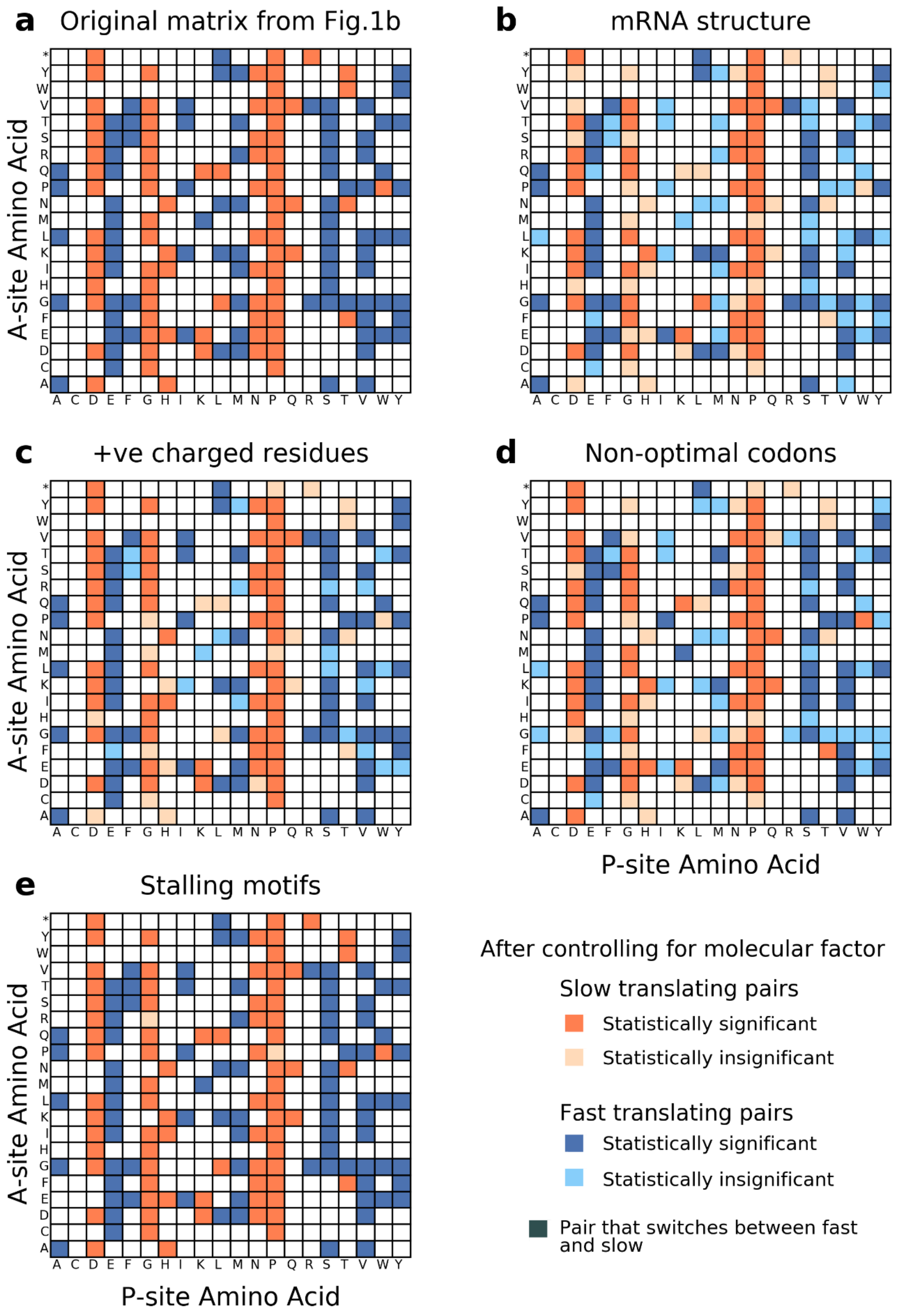
a
Original matrix from Fig.1b
b
mRNA structure
c
+ve charged residues
d
Non-optimal codons
e
Stalling motifs
A-site Amino Acid
P-site Amino Acid
* Y W V T S R Q P N M L K I H G F E D C A
A C D E F G H I K L M N P Q R S T V W Y
After controlling for molecular factor
Slow translating pairs
Statistically significant
Statistically insignificant
Fast translating pairs
Statistically significant
Statistically insignificant
Pair that switches between fast and slow

**Figure S3: The sign of the percent change in ribosome density (Eq. 1) for the fast- and slow- translating amino acid pairs remains the same after controlling for different molecular factors known to influence translation speed. (a)** 86 fast translating pairs (dark blue) and 81 slow translating pairs (dark orange) as reported in Fig. 1b. **(b)** The same 167 significant amino acid pairs after controlling for downstream mRNA secondary structure. The direction of the median speed change for all 167 pairs remains the same but 72 (43%) of pairs lose statistical significance (light orange and light blue colors for slow and fast, respectively). **(c)** Same analysis as (b) but controlling for positively charged residues 2 to 5 residues present upstream of the P-site. The direction of the speed change for all 167 pairs remains the same, 43 (26%) pairs lose statistical significance. **(d)** Same analysis as (b) but controlling for non-optimal codons in the A-site. The direction of the median speed change for all 167 pairs remains the same but 60 (36%) pairs lose statistical significance. **(e)** Similar analysis as (b) but controlling for proline containing tripeptide stalling motifs. The direction of the speed change for all 167 pairs remains the same, 2 pairs (2.4%) lose statistical significance. The loss of statistical significance is primarily due to a decrease in the sample size after leaving out of instances of the given molecular factor from the $[\rho(X,Z)]$ and $[\rho(\sim X,Z)]$ distributions that are compared.

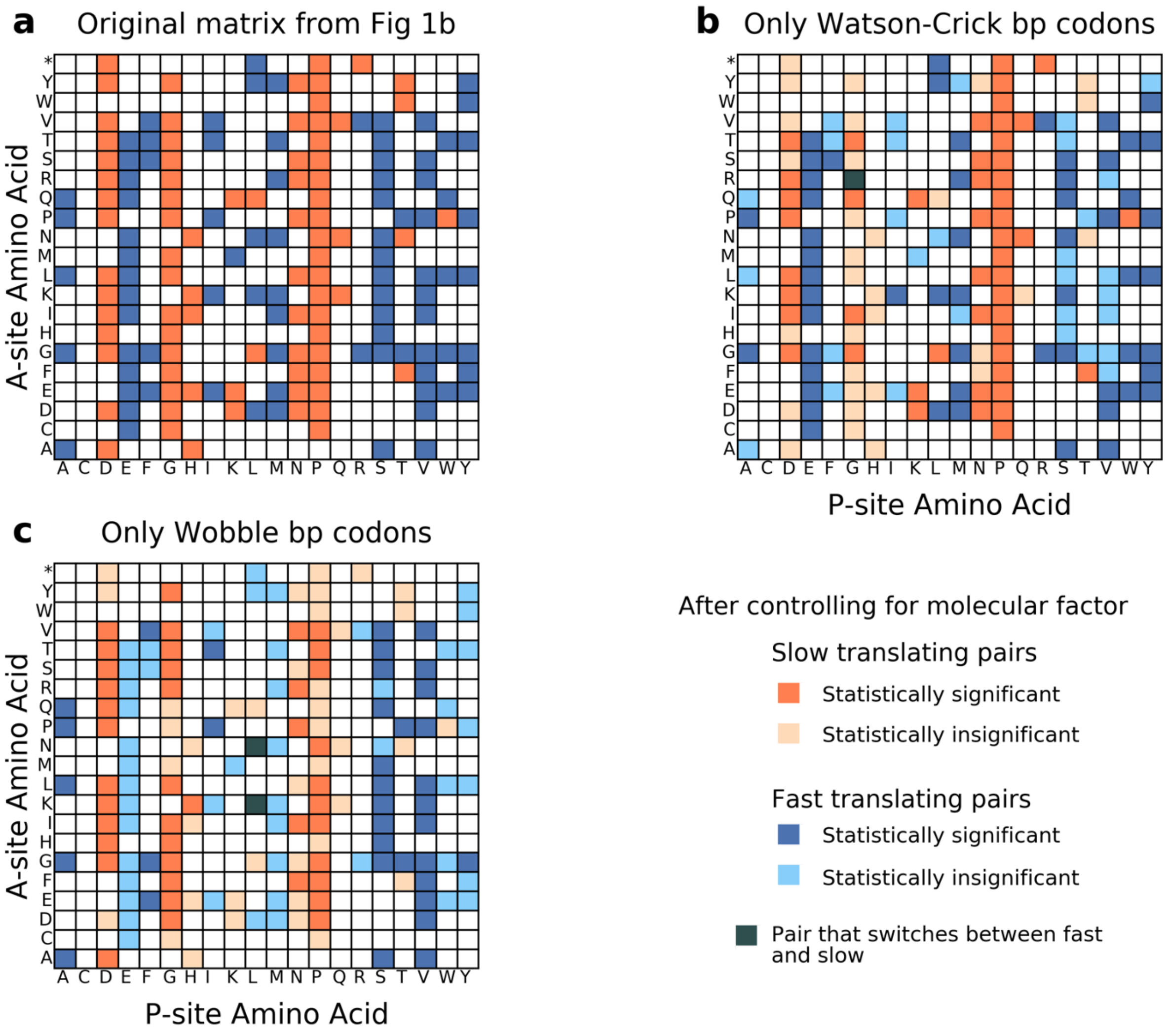


**Figure S4: Patterns of sign of the percent change of Eq. 1 are not explained by differences in Wobble decoding versus Watson Crick decoding in the P- and A-sites. (a)** 86 fast translating pairs (dark blue) and 81 slow translating pairs (dark orange) as reported in Fig. 1b. **(b)** Fields *et al.* 2016 demonstrated that codons pairs decoded by wobble base pairing mechanism may cause translational slowdown. However, if we only consider instances of amino acid pairs coming from tRNAs that decode codons through Watson-Crick geometry, the sign of the percent change in median normalized ribosome density $\rho$ (Eq. 1) remains the same in 166 out of 167 the fast and slow translating pairs identified in Fig. 1b. The coloring scheme is the same as in Fig S2. **(c)** Similarly, if we restrict our analysis to instances of amino acid pairs that are encoded by wobble base pairing tRNAs in both the P- and A-sites, the sign of the percent change in median normalized ribosome density $\rho$ (Eq. 1) remains the same in 165 out of 167 significant fast and slow translating pairs identified in Fig. 1b.

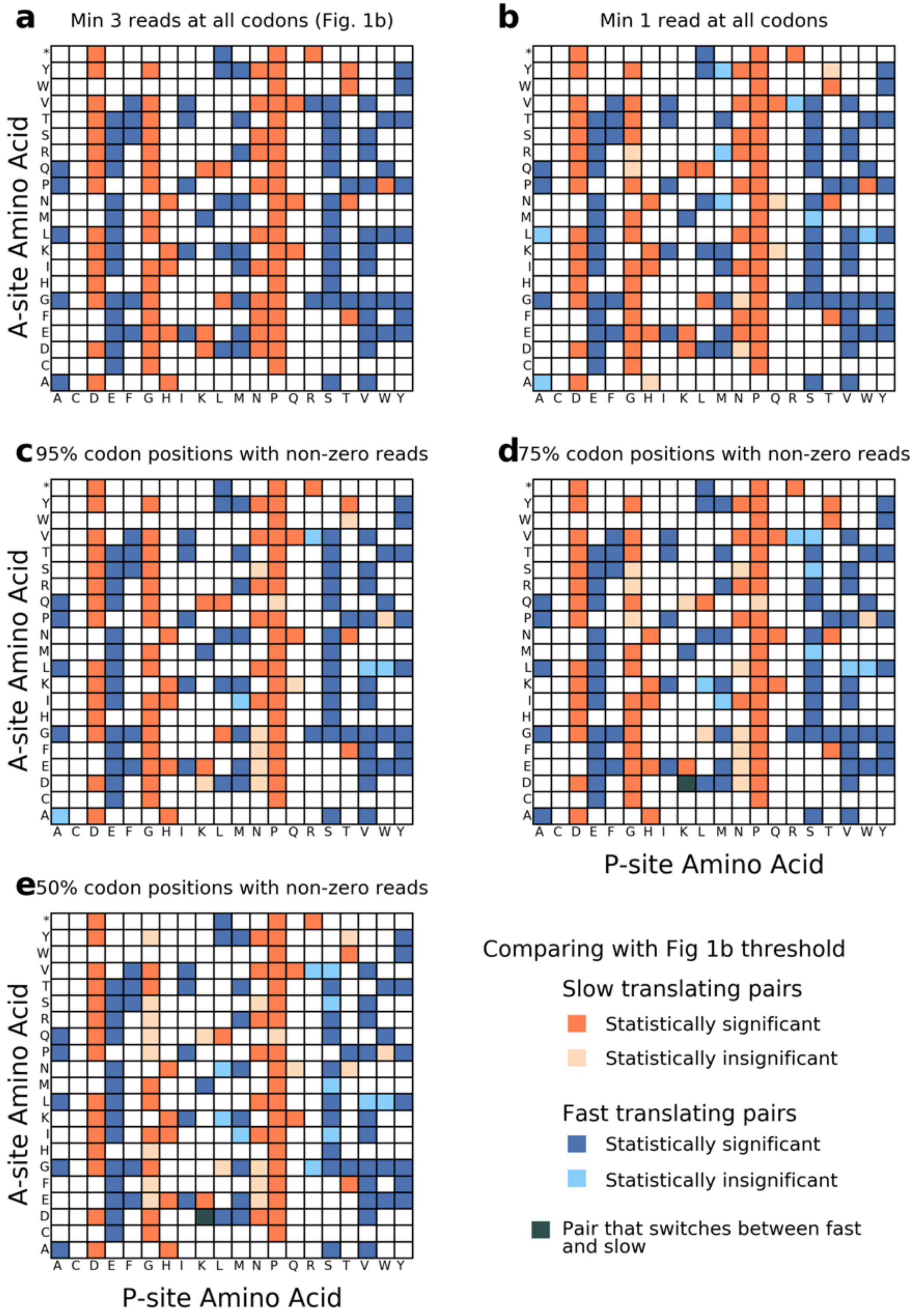
a
Min 3 reads at all codons (Fig. 1b)
b
Min 1 read at all codons
c
95% codon positions with non-zero reads
d
75% codon positions with non-zero reads
e
50% codon positions with non-zero reads
A-site Amino Acid
P-site Amino Acid
* Y W V T S R Q P N M L K I H G F E D C A
A C D E F G H I K L M N P Q R S T V W Y
Comparing with Fig 1b threshold
Slow translating pairs
Statistically significant
Statistically insignificant
Fast translating pairs
Statistically significant
Statistically insignificant
Pair that switches between fast and slow

**Figure S5: The sign of the percent change in normalized ribosome density (Eq. 1) for the fast- and slow- translating amino acid pairs remains the same at different read-depth thresholds used in the gene selection criteria.** The original matrix from Fig. 1b is represented in **(a)** where the dataset includes genes that have at least 3 reads in mapped in every codon position $(n = 364)$. When we relax the criteria to include genes that have even 1 read in every codon position $n = 687)$ **(b)** or 1 read in at least 95% of the codon position $(n = 1{,}745)$ **(c)**, we see that the sign of percent change is the same for all 167 pairs. For the selection criteria of **(d)** at least 75% of codon positions have non-zero reads $(n = 3{,}150)$ and **(e)** at least 50% of codon positions have non-zero reads $(n = 3{,}986)$, we see that the sign of percent change is same for 166-out-of-167 pairs and switches from slow to fast for (K, D) pair.

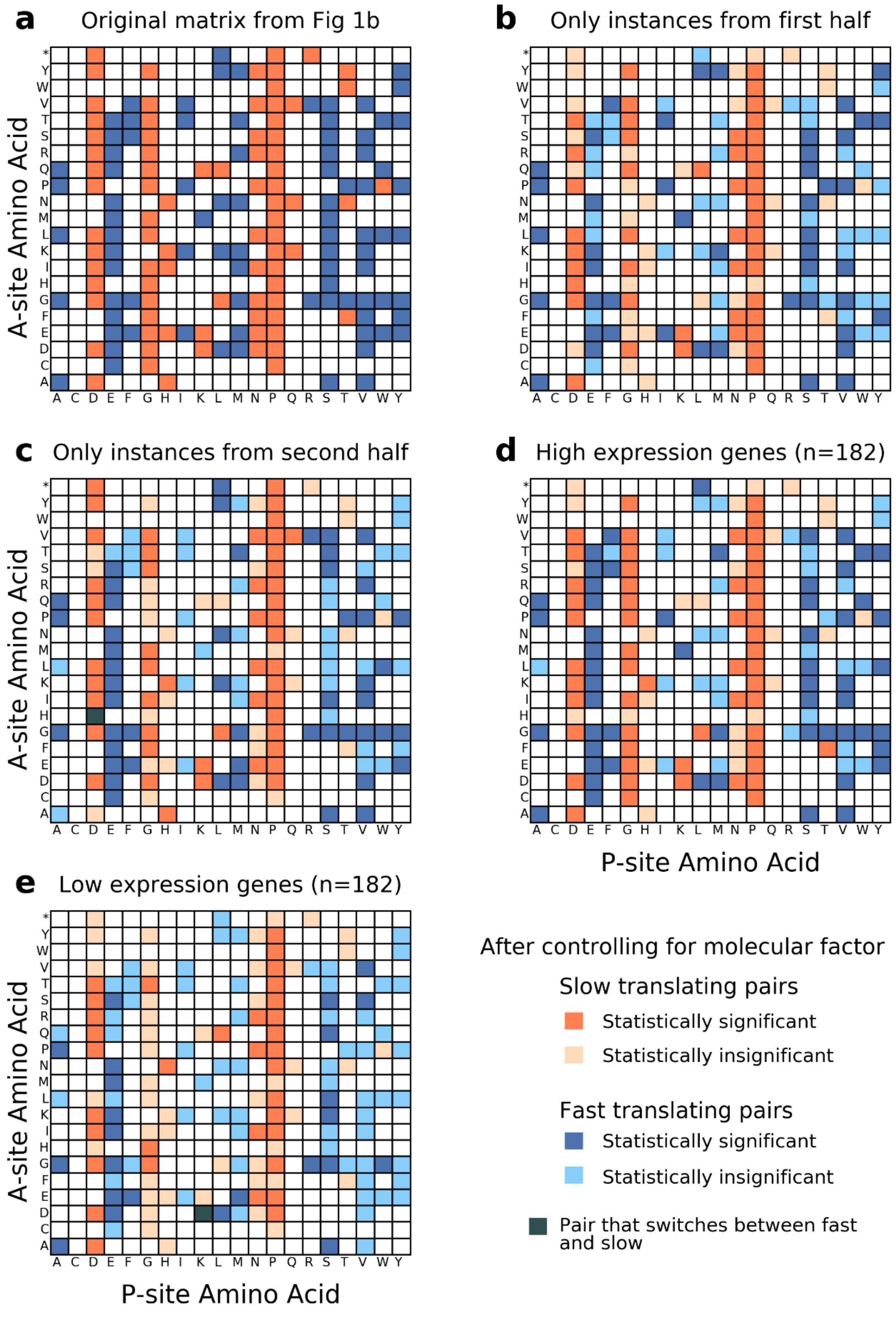

a Original matrix from Fig 1b
b Only instances from first half
c Only instances from second half
d High expression genes (n=182)
e Low expression genes (n=182)
A-site Amino Acid
P-site Amino Acid
* Y W V T S R Q P N M L K I H G F E D C A
A C D E F G H I K L M N P Q R S T V W Y
After controlling for molecular factor
Slow translating pairs
Statistically significant
Statistically insignificant
Fast translating pairs
Statistically significant
Statistically insignificant
Pair that switches between fast and slow

**Figure S6: The sign of the percent change in normalized ribosome density (Eq. 1) for the fast- and slow- translating amino acid pairs remains the same irrespective of the expression level and regions of the transcripts from where the instances of amino acid pairs are chosen. (a)** The original matrix from Fig. 1b but showing the sign change. **(b)** The matrix where the instances forming the distribution (X,Z) come from only the first half of the transcript. The direction of percent change is same for all 167 amino acid pairs. **(c)** The matrix where the instances forming the distribution (X,Z) come from only the second half of the transcript. The direction of percent change is same for 166-out-of-167 pairs. **(d)** The matrix where the instances forming the distribution (X,Z) come from only the highly expressed gene set of our data $(n = 182)$. The direction of percent change is same for all 167 amino acid pairs. **(e)** The matrix where the instances forming the distribution (X,Z) come from only low expressed gene set of our data $(n = 182)$. The direction of percent change is same for 166-out-of-167 pairs.

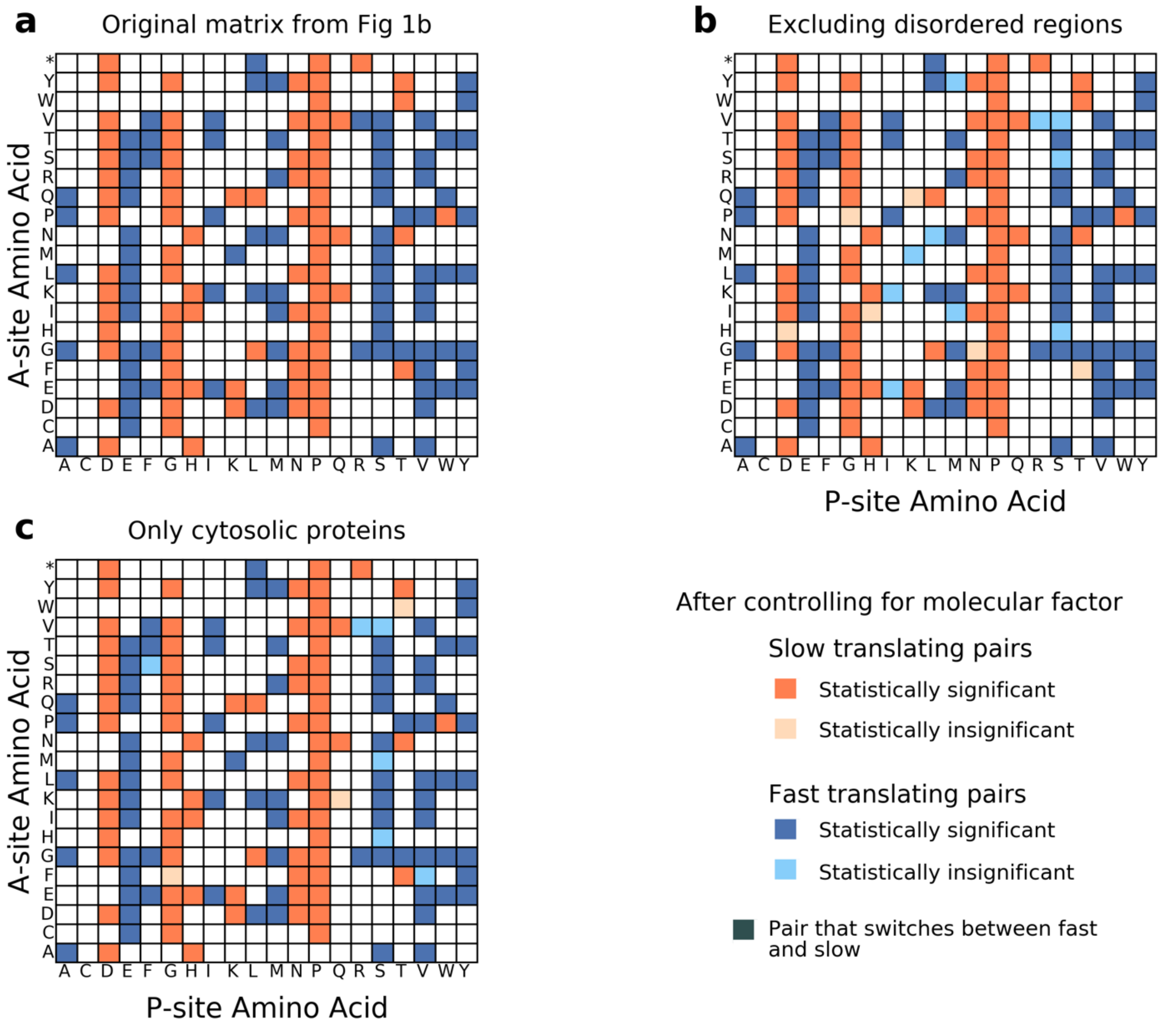


**Figure S7: The sign of the percent change in normalized ribosome density (Eq. 1) for the fast- and slow- translating amino acid pairs remains the same while controlling for regions of the transcripts from which the amino acid pairs arise. (a)** The original matrix from Fig. 1b. **(b)** The matrix where the instances forming the distribution (X,Z) that arise from disordered regions of our gene transcript dataset are excluded. The direction of percent change is same for all 167 amino acid pairs. **(c)** The matrix where the instances forming the distribution (X,Z) are chosen only from transcripts encoding cytosolic proteins that can avoid gene regions encoding signal peptides and transmembrane helices. The direction of percent change is same for all 167 pairs.

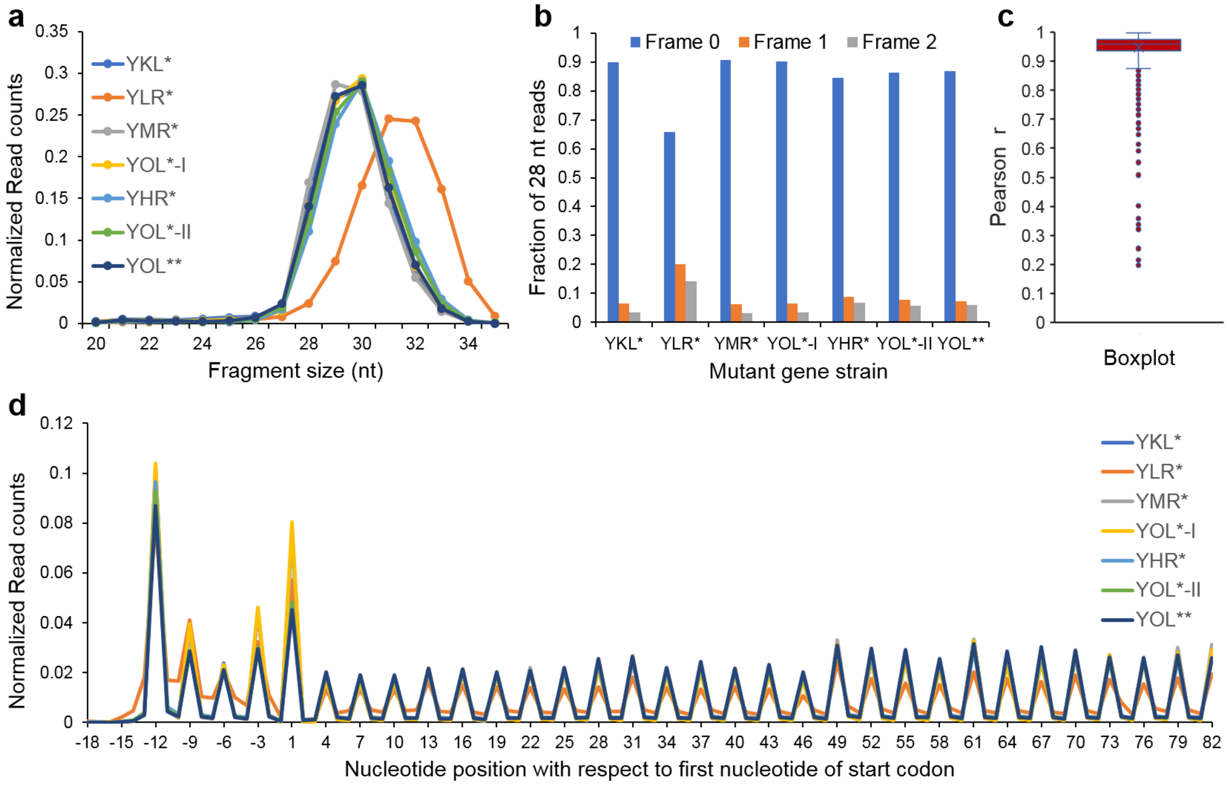


**Figure S8: The ribosome profiling data for all the mutant strains demonstrate consistent fragment size distribution, strong 3 nt periodicity, robust frame distribution and high pairwise correlation of individual transcript's ribosome profiles. (a)** Fragment size distribution of reads mapped to the CDS regions by 5′ end including 50 nt region upstream of start codon for the mutant strains created in this study. YOL* strain was subjected to ribosome profiling twice on different days denoted as YOL*-I and YOL*-II for phases I and II respectively (see Methods for details). **(b)** The distribution of reads of fragment size 28 whose 5′ end have aligned to reading frame 0, 1 or 2 for all mutant strains. **(c)** The pairwise correlation of ribosome profiles for 108 high coverage genes that have at least 3 reads at every codon position. The pairwise correlation is carried out only between samples prepared during the same phase (See Methods for details). The median Pearson r is 0.96 indicating very high correlation between ribosome profiles of genes across different samples. **(d)** Meta-gene profile of normalized read counts for fragments of size 28 mapped by the 5′ end and plotted in a 100 nt region starting from -18 nucleotide position with respect to first nucleotide of start codon up to nucleotide position 82. For analyses in (a), (b) and (d), for all mutant strains, the normalized read counts were averaged across all replicates.

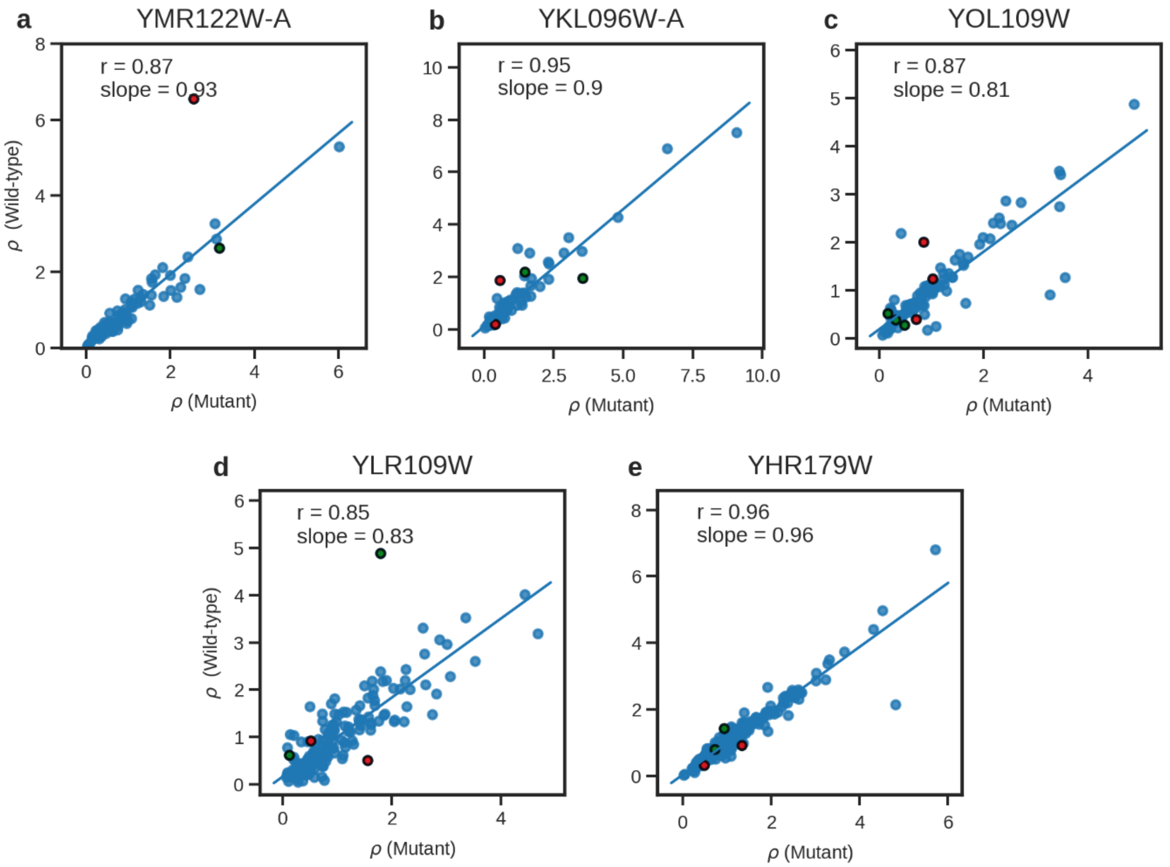


**Figure S9: Ribosome profiles of mutant and wild-type strains are highly correlated. (a-e)** The normalized ribosome density $\rho$ for each codon in the mutated gene, averaged over all replicates for mutant and wild-type reference samples are correlated and plotted on the X and Y-axis respectively. In all cases, the median Pearson R between the individual replicates is greater than 0.83. The mutated amino acids are represented by green data points while $\rho$ for codon position that is in the A-site when the mutated amino acid is in P-site is shown by red data points.

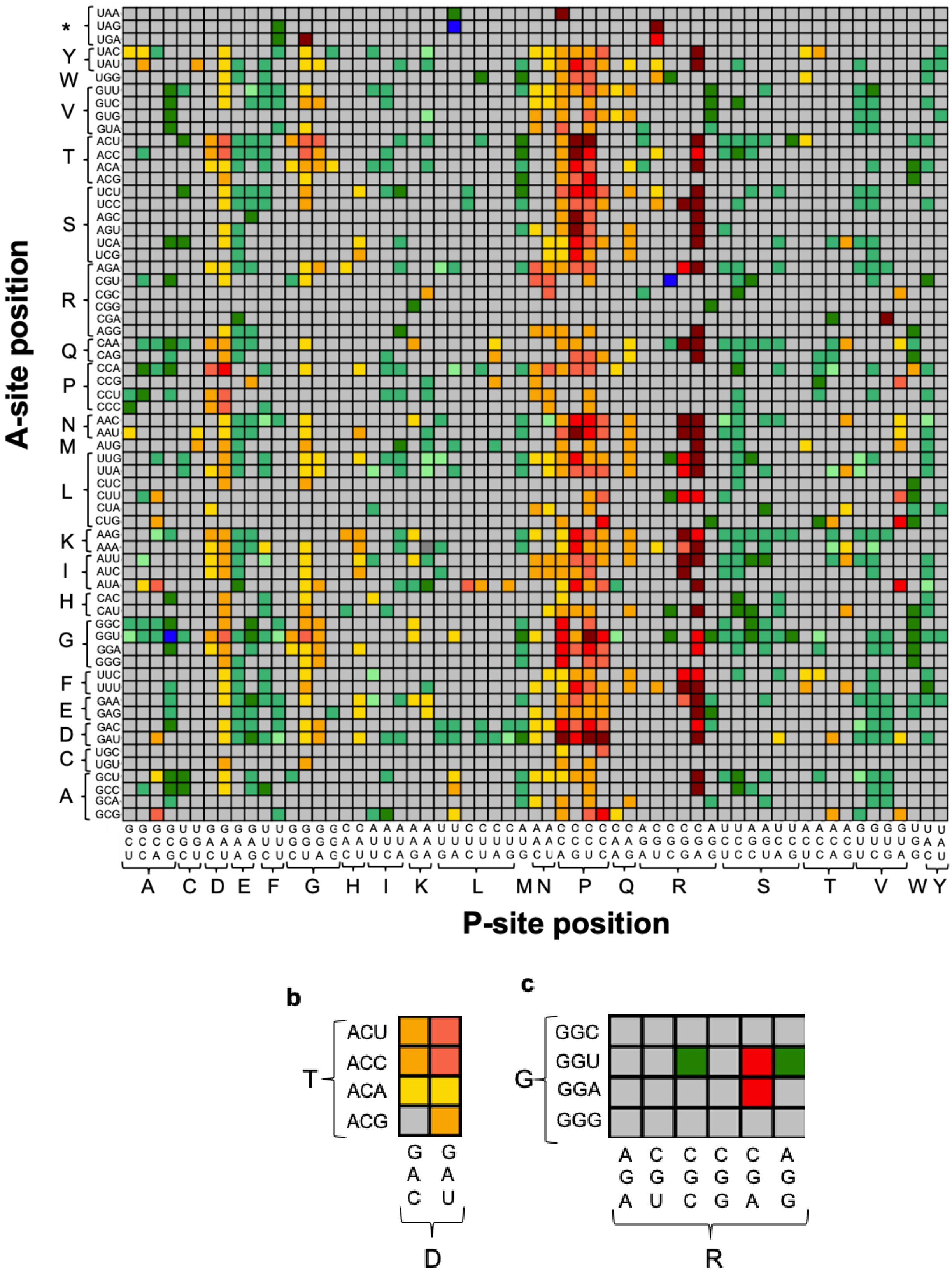
A-site position
P-site position
* Y W V T S R Q P N M L K I H G F E D C A
A C D E F G H I K L M N P Q R S T V W Y
b
T
ACU
ACC
ACA
ACG
GAC
GAU
D
c
G
GGC
GGU
GGA
GGG
AGA
CGU
CGC
CGG
CGA
AGG
R

**Figure S10: The effect of amino acid pair is qualitatively consistent among subset of codon pairs except for few cases where opposing effects are present. (a)** The 21X20 amino acid matrix in Fig. 1b is now projected to 64X61 codon level matrix. Each box in the matrix represents the pair of codons that are present in the P-site and the A-site of the ribosome and the color indicates the percent change in median normalized ribosome density $\rho$ when that particular codon is in P-site compared to any other codon in the P-site, keeping the A-site codon constant (Eq. 1). The sign of the percent change must be consistent in all 6 analyzed ribosome profiling datasets and statistically significant in at least 4 out of the 6 datasets, otherwise the box is colored gray. The subset of codon pairs for an amino acid pair is shown here for **(a)** amino acid pair (D, T) and **(b)** amino acid pair (R, G). **(a)** For the 8 possible codon pair combinations for pair (D, T), 7 of them show the same effect of statistically significant slowdown while the 8$^{th}$ codon pair is not significant. **(b)** Amino acid pair (R, G) is a fast-translating pair (Fig. 1b) but when the effect is broken down to the 24 possible codon pair combinations, 2 codon pairs show a significant speedup (green boxes) while if CGA codon of R is used in the A-site, it can result in a slowdown. This indicates for that for some amino acid pairs, the tRNA-tRNA and tRNA-mRNA interactions can show opposing effects.

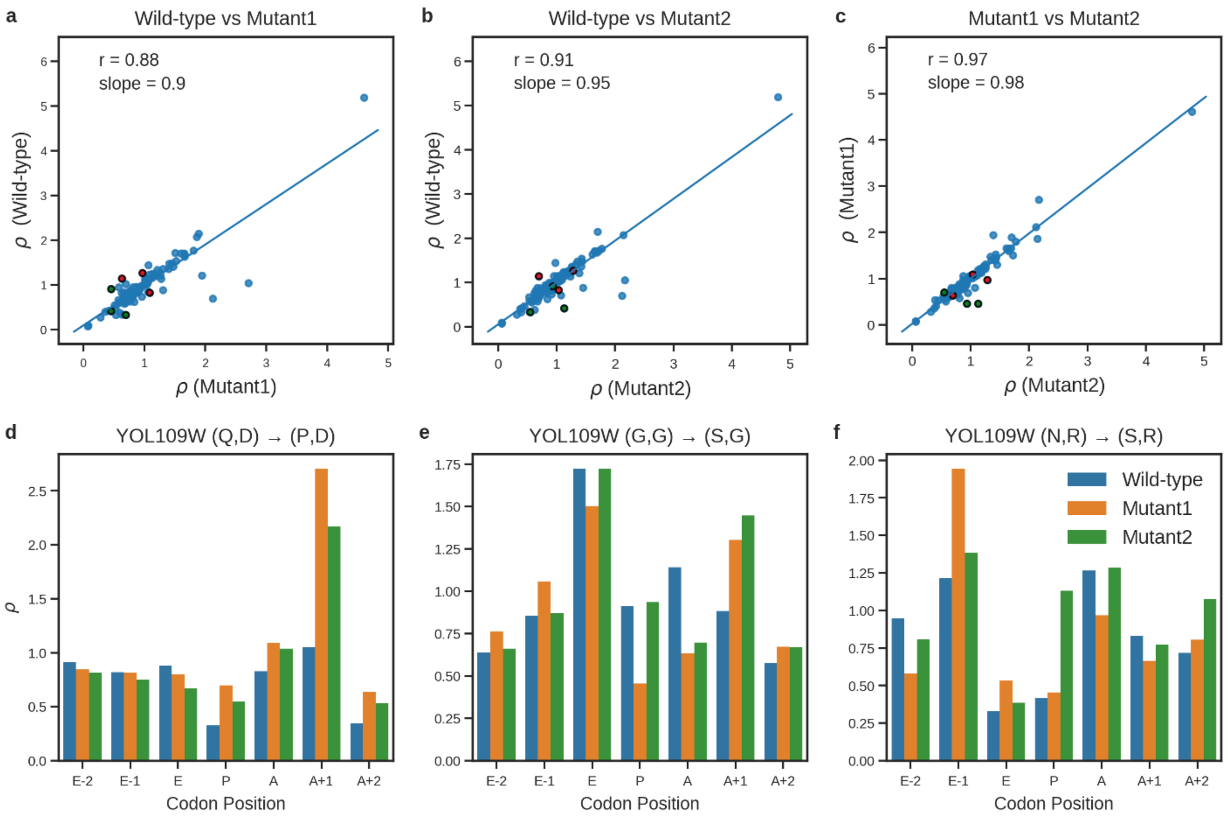


**Figure S11: Ribosome profiles are highly correlated for the two mutant strains created using synonymous codons with wild-type as well as with each other for 3 codon positions across YOL109W. (a-c)** The normalized ribosome density $\rho$ profile across the entire transcript of YOL109W is correlated for all combinations of the three strains: Wild-type, Mutant1 and Mutant2. The mutated position is shown in green while the next codon position which is in the A-site when the mutated codon position is in P-site is shown in red. **d-f)** For the three mutations, the normalized ribosome density $\rho$ is shown for the region around the mutated codon with the mutated position in the P-site. The change in $\rho$ for the A-site position is represented in Fig. 3c-e.

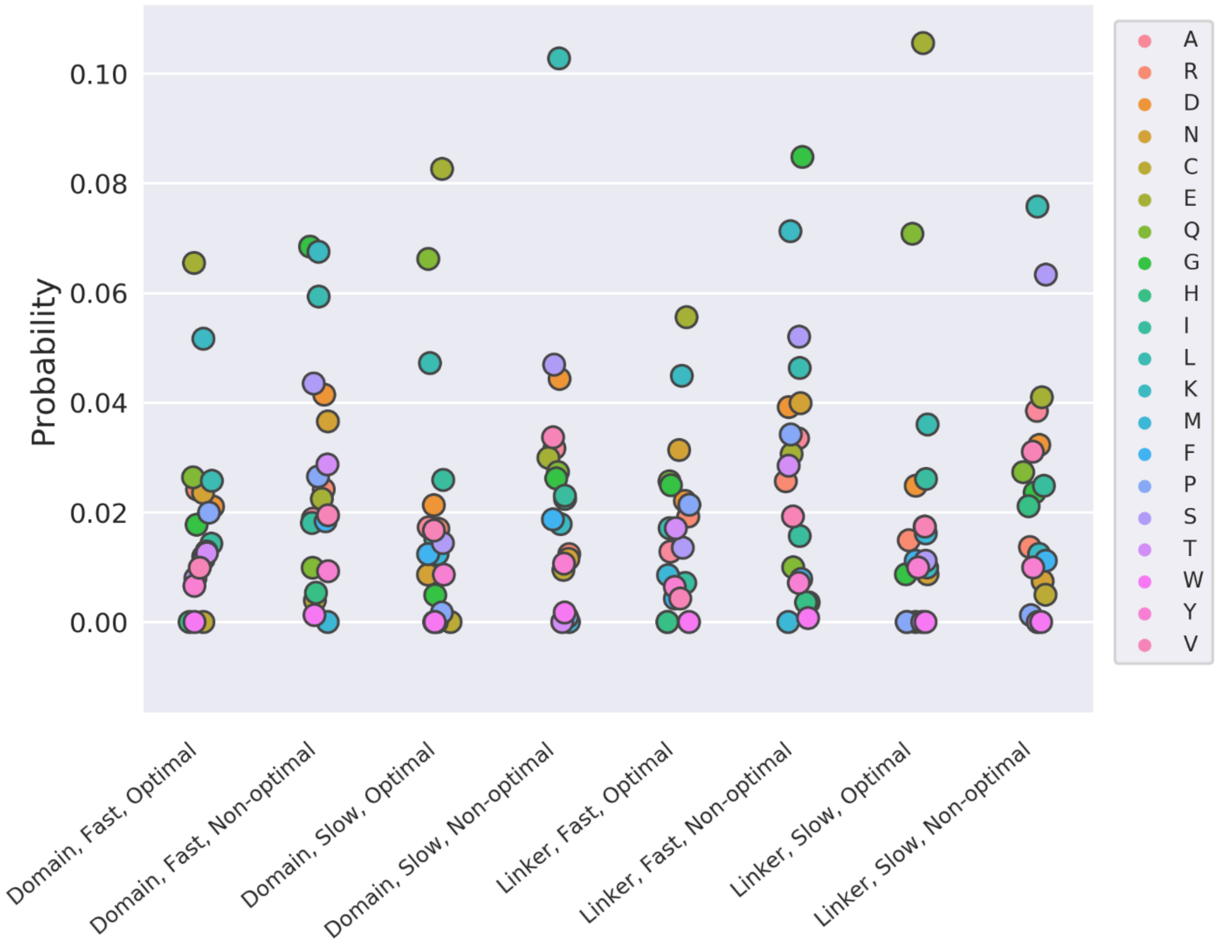


**Figure S12: Optimal and non-optimal codons are equally distributed between the domain and linker regions of proteins for both fast- and slow-translating amino acid pairs**. Probability of observing optimal or non-optimal codons in either domain or linker regions, and whether the codons are part of fast- or slow-translating amino acids identified in Fig. 1b. Comparison of domain versus linker regions for both optimal and non-optimal codons and also for optimal vs non-optimal codons within both domain and linker regions in slow pairs shows that they are equally distributed ($p - values > 0.05$, Wilcoxon-signed rank test, corrected for multiple testing by Bonferroni method).

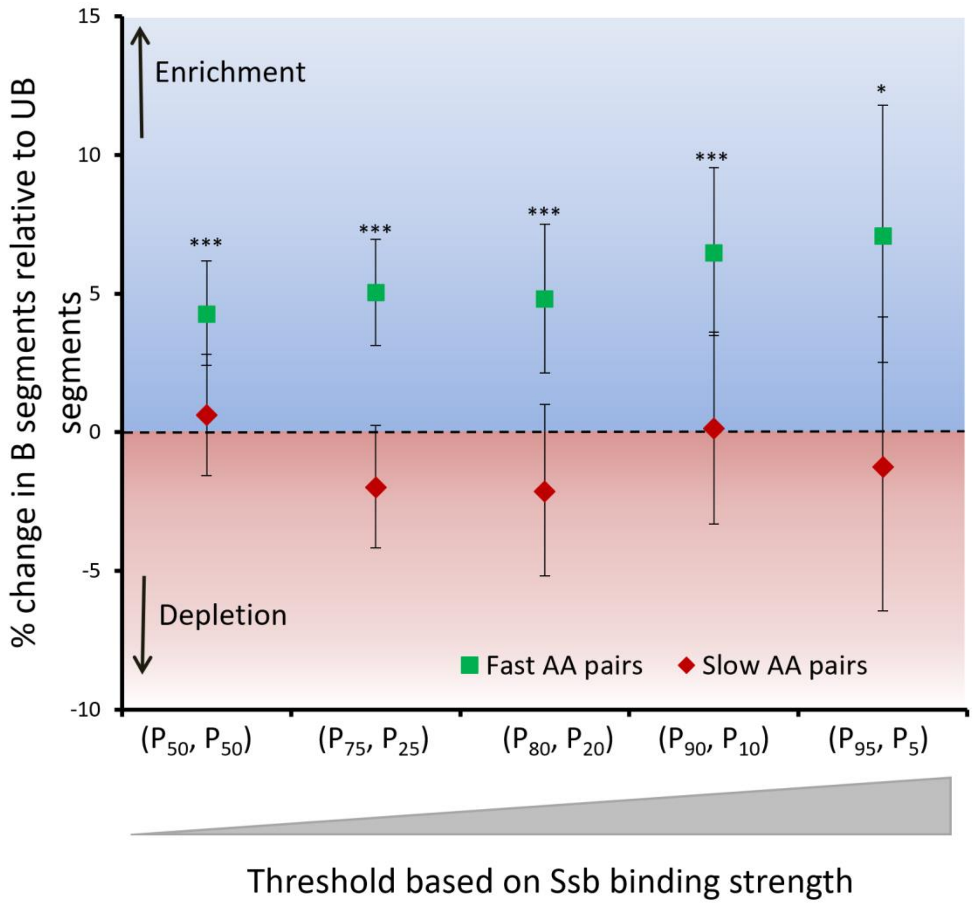


**Figure S13: Fast-translating amino acid pairs are enriched in those transcript segments that are being translated when the chaperone Ssb is bound to the nascent chain**. As done in a previous publication [3], 5 different thresholds listed on the x-axis were defined based on the percentile values of the Cumulative Distribution Function of Fold Enrichment (FE) metric (see Methods for details). For each of these thresholds, Ssb-bound translating regions (B segments) were defined by the nucleotide positions that have FE values greater than the upper threshold (*e.g.*, $P_{75}$ for threshold ($P_{75}$, $P_{25}$)) while Ssb-unbound translating regions (UB segments) are defined by the nucleotide positions with FE less than the lower threshold (*e.g.*, $P_{25}$ for threshold ($P_{75}$, $P_{25}$)). For fast-translating amino acid pairs (green) and slow-translating amino acid pairs (red), the percent change in probability of finding these pairs in the B segments relative to UB segments is reported on the y-axis. A positive percent change indicates enrichment and negative percent change indicates depletion of the pairs. Significance of enrichment/depletion is calculated using the random permutation test (***: $p < 0.0001$, **: $p < 0.01$, *: $p < 0.05$). Error bars represent 95% CI and were estimated using the bootstrapping method. These results indicate that the fast translating pairs of amino acids are enriched in those segments being translated when Ssb is bound to the ribosome nascent chain complex.

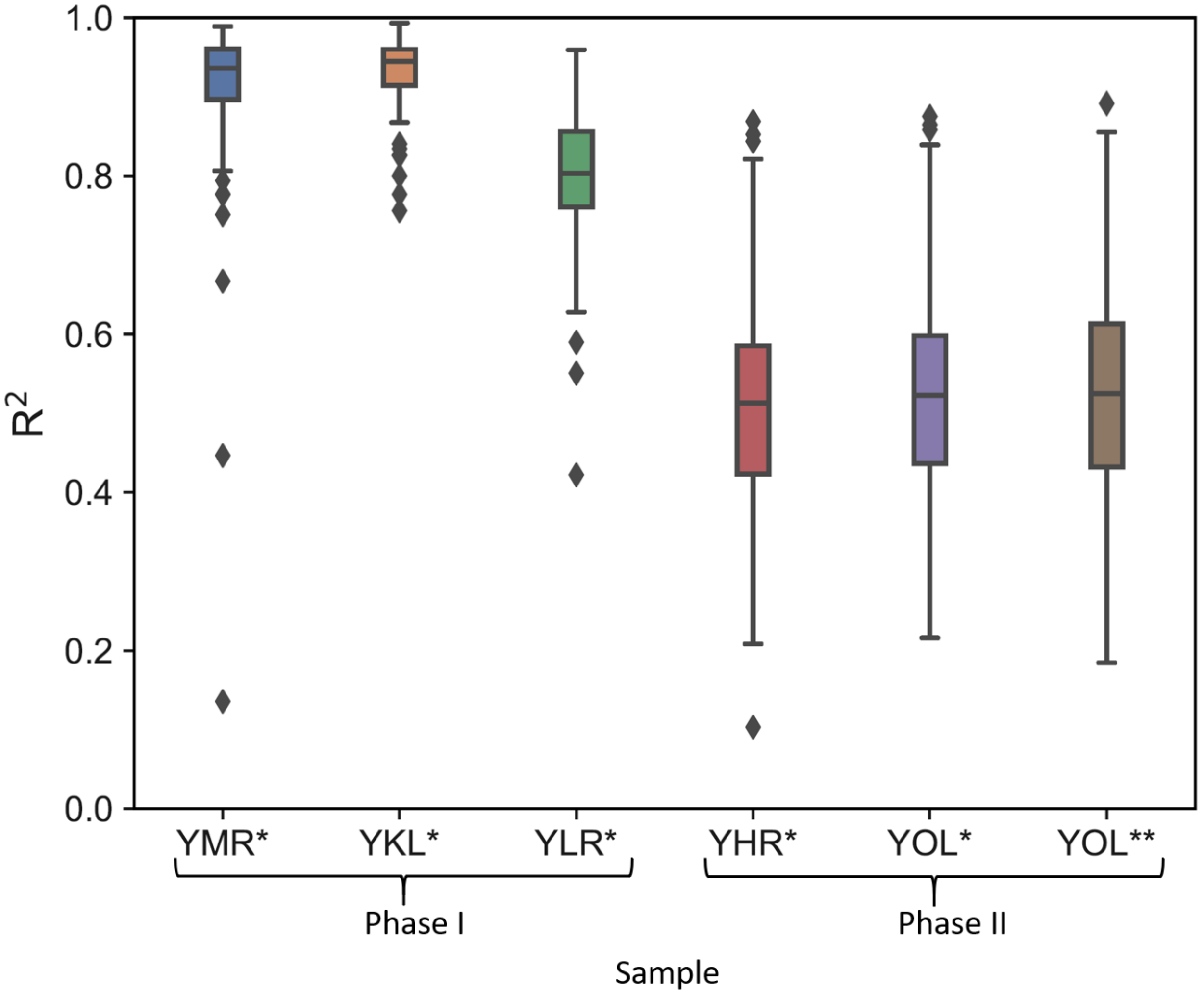


**Figure S14: Samples prepared in the same phase (single batch on same day) exhibit higher correlations than samples prepared in different phases.** For 118 genes having at least 3 reads per codon in the highest coverage replicate of YOL* mutant strain prepared in Phase I, pairwise correlations are run between the normalized ribosome densities across the CDS in this YOL* replicate with the highest coverage replicate of all other mutant strains. The boxplot of $R^2$ values is plotted here for correlating the normalized ribosome density profiles of 118 genes of sample YOL* from Phase I with mutant strains from Phase I and II. YOL* mutant strain prepared in Phase I has highly correlated ribosome densities at individual codon positions with mutant strains YMR*, YKL* and YLR* which were also prepared in Phase I. The correlation of codon level ribosome density for 118 YOL* genes was lower when correlated with ribosome profiles of strains YHR*, YOL* and YOL** prepared in Phase II. Hence, wild-type replicates were chosen for our mutations (Fig. 2) such that they have been prepared in the same phase.

**Table S1: Ribosome profiling data was obtained from five different published studies. The study and the sample accession numbers are listed below.**

| Dataset (first author name) | Year of publication | Number of replicates | GEO Study | Accession numbers of samples used |
|---|---|---|---|---|
| Jan [6] | 2014 | 1 | GSE61012 | GSM1495525 |
| Williams [4] | 2014 | 1 | GSE61011 | GSM1495503 |
| Young [8] | 2015 | 1 | GSE69414 | GSM1700885 |
| Weinberg [7] | 2016 | 1 | GSE53268 | GSM1289257 |
| Nissley [5] | 2016 | 2 | GSE75322 | GSM1949550, GSM1949551 |

**Table S2: Details on the 12 single amino acid mutations that were made across 5 different genes.** For a given amino acid pair in the A- and P-sites, the columns, from left to right, report the amino acid in the P-site (‘P-site’), the amino acid that the P-site is mutated to (‘Mutated P-site’), the amino acid in the A-site (‘A-site’), the Gene name (‘Gene’), the A-site codon number (‘A-site codon No.’), the wild-type and mutated codon in the P-site (P-site codon and Mutated P-site codon, respectively), and the codon in the A-site (‘A-site codon’).

| P-site | Mutated P-site | A-site | Gene | A-site Codon No. | P-site Codon | Mutated P-site codon | A-site Codon |
|---|---|---|---|---|---|---|---|
| Slow → Fast translation mutations | | | | | | | |
| P | E | G | YMR122W-A | 56 | GGC | GAA | CCA |
| N | S | R | YOL109W | 106 | AAC | UCC | CGU |
| D | F | G | YKL096W-A | 32 | GGU | UUC | GAC |
| G | S | T | YLR109W | 162 | ACC | UCU | GGU |
| G | S | G | YOL109W | 99 | GGC | UCU | GGU |
| Fast → Slow translation mutations | | | | | | | |
| Q | P | D | YOL109W | 14 | CAA | CCA | GAU |
| S | G | G | YLR109W | 140 | GGU | GGU | AGU |
| S | G | T | YKL096W-A | 62 | ACC | GGU | AGC |
| V | H | K | YHR179W | 339 | GUG | CAC | AAG |
| E | K | E | YHR179W | 150 | GAA | AAA | GAA |
| Negative control mutations | | | | | | | |
| V | Y | F | YHR179W | 251 | GUC | UAC | UUC |
| L | N | A | YHR179W | 129 | CUU | AAC | GCU |

**Table S3: Statistics of read mapping for ribosome profiling experiments for the mutant strains carried out in this study**. 1.7 billion reads were mapped to the exome in total for all samples with an average of 86 million reads per sample.

| | Sample | Total reads (millions) | Mapped to rRNA | | Mapped to exome | |
|---|---|---|---|---|---|---|
| | | | Reads (millions) | % reads | Reads (millions) | % reads |
| **Phase I samples** | YMR* rep1 | 37.10 | 9.04 | 24.37% | 24.76 | 66.75% |
| | YMR* rep2 | 56.06 | 11.90 | 21.23% | 25.02 | 44.63% |
| | YKL* rep1 | 59.79 | 15.65 | 26.18% | 41.46 | 69.35% |
| | YKL* rep2 | 65.39 | 17.04 | 26.05% | 44.88 | 68.64% |
| | YOL* rep1 | 123.58 | 28.99 | 23.46% | 83.56 | 67.62% |
| | YOL* rep2 | 59.10 | 14.76 | 24.97% | 41.51 | 70.25% |
| | YLR* rep1 | 55.39 | 8.54 | 15.43% | 44.03 | 79.49% |
| | YLR* rep2 | 57.00 | 5.34 | 9.36% | 44.67 | 78.36% |
| **Phase II samples** | YHR* rep1 | 195.94 | 27.96 | 14.27% | 158.14 | 80.71% |
| | YHR* rep2 | 163.22 | 30.91 | 18.94% | 124.51 | 76.28% |
| | YHR* rep3 | 151.04 | 24.84 | 16.45% | 116.16 | 76.91% |
| | YHR* rep4 | 159.08 | 19.18 | 12.06% | 131.19 | 82.46% |
| | YOL* rep1 | 158.03 | 29.91 | 18.93% | 118.97 | 75.28% |
| | YOL* rep2 | 148.15 | 16.85 | 11.38% | 99.34 | 67.05% |
| | YOL* rep3 | 149.12 | 24.39 | 16.36% | 114.79 | 76.98% |
| | YOL* rep4 | 141.42 | 23.34 | 16.50% | 104.26 | 73.72% |
| | YOL** rep1 | 146.85 | 18.53 | 12.62% | 74.68 | 50.85% |
| | YOL** rep2 | 157.73 | 22.76 | 14.43% | 125.87 | 79.80% |
| | YOL** rep3 | 141.19 | 20.74 | 14.69% | 112.96 | 80.00% |
| | YOL** rep4 | 141.42 | 23.34 | 16.50% | 104.26 | 73.72% |
| | | | | **Average** | 86.75 | |
| | | | | **Total reads mapped to exome** | 1735.10 | |

**Table S4: Three mutations in gene YOL109W to test the contribution of amino acid and tRNA identity.** Columns are the same as in Table S2, except the two synonymous mutations that encode for the same mutated residue at the P-site, are labeled 'Mutant 1 P-site codon' and 'Mutant 2 P-site codon'.

| P-site | Mutated P-site | A-site | Gene | A-site Codon no | P-site Codon | Mutant 1 P-site codon | Mutant 2 P-site codon | A-site Codon |
|---|---|---|---|---|---|---|---|---|
| G | S | G | YOL109W | 99 | GGC | UCU | AGC | GGU |
| Q | P | D | YOL109W | 14 | CAA | CCA | CCU | GAU |
| N | S | R | YOL109W | 106 | AAC | UCC | UCG | CGU |

**Table S5: 2X2 contingency table for calculation of odds ratio and determining the enrichment/depletion of significant amino acid pairs across the proteome.**

| | Fast-translating pairs | Slow-translating pairs |
|---|---|---|
| Enriched | 18 | 12 |
| Depleted | 8 | 23 |

**Data S1: List of 364 high-coverage transcript whose ribosome profiling data is used for bioinformatic analyses in this study (Excel Spreadsheet).**

**Data S2: For all possible 7,980 mutations that are possible for the P-site residue in a P- and A-site amino acid pair, the statistical significance and the odds ratio between the two normalized ribosome density distributions are listed (Excel Spreadsheet).**

**Data S3: The sample size, percent change and statistical significance for the 64X61 matrix for the codon pairs in the P- and A-sites shown in Fig. S10. (Excel Spreadsheet).**

**Data S4: Database of 864 proteins in *S. cerevisiae* with annotated domain regions.** The file contains a header describing the terms for each protein entry and definition of terms for each annotated domain of the protein. **(Text file).**

**Supplementary references:**

[1] J. Cherry, E. Hong, C. Amundsen, R. Balakrishnan, G. Binkley, C. ET, C. KR, C. MC, D. SS, E. SR, F. DG, H. JE, H. BC, K. K, K. CJ, M. SR, N. RS, P. J, S. MS, S. M, W. S, W. ED, Saccharomyces Genome Database: the genomics resource of budding yeast., Nucleic Acids Res. 40 (2012) D700-5.

[2] C. Janke, M.M. Magiera, N. Rathfelder, C. Taxis, S. Reber, H. Maekawa, A. Moreno-Borchart, G. Doenges, E. Schwob, E. Schiebel, M. Knop, A versatile toolbox for PCR-based tagging of yeast genes: new fluorescent proteins, more markers and promoter substitution cassettes., Yeast. 21 (2004) 947–62. https://doi.org/10.1002/yea.1142.

[3] K. Döring, N. Ahmed, T. Riemer, H.G. Suresh, Y. Vainshtein, M. Habich, J. Riemer, M.P. Mayer, E.P. O’Brien, G. Kramer, B. Bukau, Profiling Ssb-Nascent Chain Interactions Reveals Principles of Hsp70-Assisted Folding, Cell. 170 (2017) 298-311.e20. https://doi.org/10.1016/j.cell.2017.06.038.

[4] C.C. Williams, C.H. Jan, J.S. Weissman, Targeting and plasticity of mitochondrial proteins revealed by proximity-specific ribosome profiling, Science. 346 (2014) 748–751. https://doi.org/10.1126/science.1257522.

[5] D.A. Nissley, A.K. Sharma, N. Ahmed, U.A. Friedrich, G. Kramer, B. Bukau, E.P. O’Brien, Accurate prediction of cellular co-translational folding indicates proteins can switch from post- to co-translational folding, Nat. Commun. 7 (2016) 10341. https://doi.org/10.1038/ncomms10341.

[6] C.H. Jan, C.C. Williams, J.S. Weissman, “Principles of ER cotranslational translocation revealed by proximity-specific ribosome profiling,” Science. 346 (2014) 748–751. https://doi.org/10.1126/science.1257521.

[7] D.E. Weinberg, P. Shah, S.W. Eichhorn, J.A. Hussmann, J.B. Plotkin, D.P. Bartel, Improved Ribosome-Footprint and mRNA Measurements Provide Insights into Dynamics and Regulation of Yeast Translation, Cell Rep. 14 (2016) 1787–1799. https://doi.org/10.1016/j.celrep.2016.01.043.

[8] D.J. Young, N.R. Guydosh, F. Zhang, A.G. Hinnebusch, R. Green, Rli1/ABCE1 Recycles Terminating Ribosomes and Controls Translation Reinitiation in 3′UTRs In Vivo, Cell. 162 (2015) 872–884. https://doi.org/10.1016/j.cell.2015.07.041.

[9] M. Martin, Cutadapt removes adapter sequences from high-throughput sequencing reads, EMBnet.Journal. 17 (2011) 10. https://doi.org/10.14806/ej.17.1.200.

[10] B. Langmead, S.L. Salzberg, Fast gapped-read alignment with Bowtie 2, Nat Methods. 9 (2012) 357–359. https://doi.org/10.1038/nmeth.1923.

[11] D. Kim, G. Pertea, C. Trapnell, H. Pimentel, R. Kelley, S.L. Salzberg, TopHat2: accurate alignment of transcriptomes in the presence of insertions, deletions and gene fusions, Genome Biol. 14 (2013) R36. https://doi.org/10.1186/gb-2013-14-4-r36.

[12] N. Ahmed, P. Sormanni, P. Ciryam, M. Vendruscolo, C.M. Dobson, E.P. O’Brien, Identifying A- and P-site locations on ribosome-protected mRNA fragments using Integer Programming, Sci. Reports. 9 (2019) 6256. https://doi.org/10.1038/s41598-019-42348-x.

[13] S. Rouskin, M. Zubradt, S. Washietl, M. Kellis, J.S. Weissman, Genome-wide probing of RNA structure reveals active unfolding of mRNA structures in vivo., Nature. 505 (2014)

701–705. https://doi.org/10.1038/nature12894.

[14] S. Pechmann, J. Frydman, Evolutionary conservation of codon optimality reveals hidden signatures of cotranslational folding., Nat. Struct. Mol. Biol. 20 (2013) 237–43. https://doi.org/10.1038/nsmb.2466.

[15] A.P. Schuller, C.C.C. Wu, T.E. Dever, A.R. Buskirk, R. Green, eIF5A Functions Globally in Translation Elongation and Termination, Mol. Cell. 66 (2017) 194-205.e5. https://doi.org/10.1016/j.molcel.2017.03.003.

[16] J.R. Iben, R.J. Maraia, tRNAomics: tRNA gene copy number variation and codon use provide bioinformatic evidence of a new anticodon:codon wobble pair in a eukaryote., RNA. 18 (2012) 1358–1372. https://doi.org/10.1261/rna.032151.111.

[17] R. van der Lee, B. Lang, K. Kruse, J. Gsponer, N. Sánchez de Groot, M.A. Huynen, A. Matouschek, M. Fuxreiter, M.M. Babu, Intrinsically disordered segments affect protein half-life in the cell and during evolution., Cell Rep. 8 (2014) 1832–1844. https://doi.org/10.1016/j.celrep.2014.07.055.

[18] P. Ciryam, R.I. Morimoto, M. Vendruscolo, C.M. Dobson, E.P. O'Brien, In vivo translation rates can substantially delay the cotranslational folding of the Escherichia coli cytosolic proteome., Proc. Natl. Acad. Sci. U. S. A. 110 (2013) E132-40. https://doi.org/10.1073/pnas.1213624110.

[19] P. Good, Permutation, Parametric, and Bootstrap Tests of Hypothesis, Third, Springer Series in Statistics, 2005. https://doi.org/10.1007/978-0-387-98135-2.
